# Single Photon smFRET. I. Theory and Conceptual Basis

**DOI:** 10.1101/2022.07.20.500887

**Authors:** Ayush Saurabh, Mohamadreza Fazel, Matthew Safar, Ioannis Sgouralis, Steve Pressé

## Abstract

We present a unified conceptual framework and the associated software package for single molecule Förster Resonance Energy Transfer (smFRET) analysis from single photon arrivals leveraging Bayesian nonparametrics, BNP-FRET. This unified framework addresses the following key physical complexities of a single photon smFRET experiment, including: 1) fluorophore photophysics; 2) continuous time kinetics of the labeled system with large timescale separations between photophysical phenomena such as excited photophysical state lifetimes and events such as transition between system states; 3) unavoidable detector artefacts; 4) background emissions; 5) unknown number of system states; and 6) both continuous and pulsed illumination. These physical features necessarily demand a novel framework that extends beyond existing tools. In particular, the theory naturally brings us to a hidden Markov model (HMM) with a second order structure and Bayesian nonparametrics (BNP) on account of items 1, 2 and 5 on the list. In the second and third companion manuscripts, we discuss the direct effects of these key complexities on the inference of parameters for continuous and pulsed illumination, respectively.

**Why It Matters:** smFRET is a widely used technique for studying kinetics of molecular complexes. However, until now, smFRET data analysis methods required specifying *a priori* the dimensionality of the underlying physical model (the exact number of kinetic parameters). Such approaches are inherently limiting given the typically unknown number of physical configurations a molecular complex may assume. The methods presented here eliminate this requirement and allow estimating the physical model itself along with kinetic parameters, while incorporating all sources of noise in the data.

## 1 Introduction

Förster Resonance Energy Transfer (FRET) has served as a spectroscopic ruler to study motion at the nanometer scale [1–4], and has revealed insight into intra- and intermolecular dynamics of proteins [5–11], nucleic acids [12], and their interactions [13, 14]. In particular, single molecule FRET (smFRET) experiments have been used to determine the pore size and opening mechanism of ion channels sensitive to mechanical stress in the membrane [15], the intermediate stages of protein folding [16, 17], and the chromatin interactions modulated by the helper protein HP1*α* involved in allowing genetic transcription for tightly packed chromatin [18].

A typical FRET experiment involves labeling molecules of interest with donor and acceptor dyes such that the donor may transfer energy to the acceptor via dipole-dipole interaction when separated by distances of 2-10 nm [19]. This interaction weakens rapidly with increasing separation *R* and goes as *R*^-6^ [20, 21].

To induce FRET during experiments, the donor is illuminated by a continuous or pulsating light source for the desired time period or until the dyes photobleach. Upon excitation, the donor may emit a photon itself or transfer its energy nonradiatively to the acceptor which eventually relaxes to emit a photon of a different color [20, 21]. As such, the data collected consists of photon arrival times (for single photon experiments) or, otherwise, brightness values in addition to photon colors collected in different detection channels.

The distance dependence in the rate of energy transfer between donor and acceptor is key in using smFRET as a molecular ruler. Furthermore, this distance dependence directly manifests itself in the form of higher fraction of photons detected in the acceptor channel when the dyes are closer together (as demonstrated in Fig. 1). This fraction is commonly referred to as the FRET efficiency,

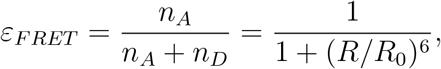

where *n_D_* and *n_A_* are the number of donor and acceptor photons detected in a given time period, respectively. Additionally, *R*_0_ is the characteristic separation that corresponds to a FRET efficiency of 0.5 or 50% of the emitted photons emanating from the acceptor.

**Figure 1:**
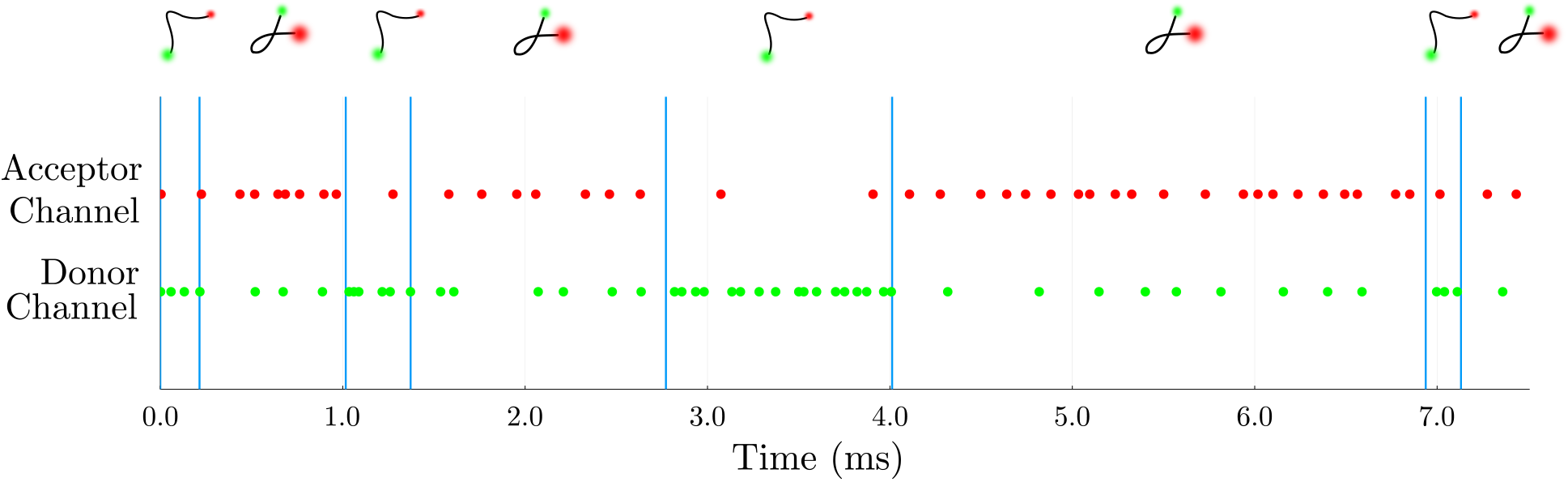
A cartoon figure illustrating smFRET data. For the experiments considered here, the kinetics along the reaction coordinate defined along the donor-acceptor distance are monitored using single photon arrival data. In the figure above, photon arrivals are represented by green dots for photons arriving into the donor channel and red dots for photons arriving in the acceptor channel. For the case where donor and acceptor label one molecule, a molecule’s transitions between system states (coinciding with conformations) is reflected by the distance between labels measured by variations in detected photon arrival times and colors.

Now, the aim of smFRET is to capture on-the-fly changes in donor-acceptor distance. However, this is often confounded by several sources of stochasticity which unavoidably obscure direct interpretation. These include: 1) the stochasticity inherent to photon arrival times; 2) a detector’s probabilistic response to an incoming photon [22]; 3) background emissions [2]; and 4) fluorescent labels’ stochastic photophysical properties [2]. Taken together, these problems necessarily contribute to uncertainty in the number of distinct system states visited by a labeled system over an experiment’s course [23–25].

Here, we delve into greater detail into items 2 and 4. In particular, item 2 pertains to questions of crosstalk, detector efficiency, dead time, dark current, and instrument response function (IRF) introducing uncertainty in excited photophysical state lifetime assessments [22, 26, 27].

Item 4 refers to a collection of effects including limited quantum yield and variable brightness due to blinking of dyes caused by nonradiative pathways [28, 29], photobleaching or permanent deactivation of the dyes [2, 28, 29], spectral overlap between the donor and acceptor dyes which may result in direct excitation of the acceptors or leaking of photons into the incorrect channel [2, 26], or a donor-acceptor pair’s relative misalignment or positioning resulting in false signals and inaccurate characterization of the separation between labeled molecules [2, 30].

Though the goal has always remained to analyze the rawest form of data, the reality of these noise properties has traditionally led to the development of approximate binned photon analyses even when data is collected at the level of single photons across two detectors. Binning is either achieved by directly summing photon arrivals over a time period when using single photon detectors [23, 31] or by integrating intensity over a few pixels when using widefield detectors [32].

While binned data analyses can be used to determine the number and connectivity of system states [33]–by computing average FRET efficiencies over bin time windows and using them in turn to construct FRET efficiency histograms [23, 25, 31, 34–36]–they come at the cost of averaging kinetics that may exist below a time bin not otherwise easily accessible [32, 37, 38]. They also eliminate information afforded by, say, the excited photophysical state lifetime in the case of pulsed illumination.

While histogram analyses are suited to infer static molecular properties, kinetics over binned time traces have also been extracted by supplementing these techniques with a hidden Markov model (HMM) treatment [23, 25, 34–36, 39].

Using HMMs, binned analysis techniques immediately face the difficulty of an unknown number of system states visited. Therefore, they require the number of system states as an input to deduce the putative kinetics between the candidate system states.

What is more, the binned analysis’ accuracy is determined by the bin sizes where large bins may result in averaging of the kinetics. Moreover, increasing bin size may lead to estimation of an excess number of system states. This artifact arises when a system appears to artificially spend more time in the system states below the bin size [38]. To address these challenges, we must infer continuous time trajectories below the bin size through, for example, the use of Markov jump processes [32] while retaining a binned, *i.e.*, discrete measurement model.

When single photon data is available we may avoid the binning issues inherent to HMM analysis [32, 40, 41]. Doing so, also allows us to directly leverage the noise properties of detectors for single photon arrivals (*e.g.*, IRF) well calibrated at the single photon level. Moreover, we can now also incorporate information available through photophysical state lifetimes when using pulsed illumination otherwise eliminated in binning data. Incorporating all of this additional information, naturally, comes with added computational cost [37] whose burden a successful method should mitigate.

Often to help reduce computational costs, further approximations on the system kinetics are invoked such as assuming system kinetics to be much slower than FRET label excitation and relaxation rates. This approximation helps decouple photophysical and system (molecular) kinetics [16, 37, 42, 43].

What is more, as they exist, the rigor of direct photon arrival analysis methods are further compromised to help reduce computational cost by treating detector features and background as preprocessing steps [16, 37, 42, 43]. In doing so, simultaneous and self-consistent inference of kinetics and other molecular features becomes unattainable. Finally, all methods, whether relying on the analysis of binned photons or single photon arrival, suffer from the “model selection problem”. That is, the problem associated with identifying the number of system states warranted by the data. More precisely, the problem associated with propagating the uncertainty introduced by items 1-4 into a probability over the models (*i.e.*, system states). Existing methods for system state identification only provide partial reprieve.

For example, while FRET histograms identify peaks to intuit the number of system states, these peaks may provide unreliable estimates for a number of reasons: 1) fast transitions between system states may result in a blurring of otherwise distinct peaks [1] or, counterintuitively, introduce more peaks [25, 38]; 2) system states may differ primarily in kinetics but not FRET efficiency [40]; 3) detector properties and background may introduce additional features in the histograms.

To address the model selection problem, overfitting penalization criteria (such as the Bayesian information criterion or BIC) [23, 44] or variational Bayesian [24] approaches have been employed.

Often, these model selection methods assume implicit properties of the system. For example, the BIC requires the assumption of weak independence between measurements (*i.e.*, ideally independent identically distributed measurements and thus no Markov kinetics in state space) and a unique likelihood maximum, both of which are violated in smFRET data [24]. Furthermore, BIC and other such methods provide point estimates rather than full probabilities over system states ignoring uncertainty from items 1-4 propagated over models [45].

As such, we need to learn distributions over system states and kinetics warranted by the data and whose breadth is dictated by the sources of uncertainty discussed above. More specifically, to address model selection and build joint distributions over system states and their kinetics, we treat the number of system states as a random variable just as the current community treats smFRET kinetic rates as random variables [25, 40, 41]. Our objective is therefore to obtain distributions over all unknowns (including system states and kinetics) while accounting for items 1-4. Furthermore, this must be achieved in a computationally efficient way avoiding, altogether, the draconian assumptions of existing in single photon analysis methods. In other words, we want to do more (by learning joint distributions over the number of system states alongside everything else) and we want it to cost less.

If we insist on learning distributions over unknowns, then it is convenient to operate within a Bayesian paradigm. Also, if the model (*i.e.*, the number of system states) is unknown, then we must further generalize to the Bayesian nonparametric (BNP) paradigm [25, 41, 46–53]. BNPs directly address the model selection problem concurrently and self-consistently while learning the associated model’s parameters and output full distributions over the number of system states and the other parameters.

In this series of three companion manuscripts, we present a complete description of single photon smFRET analysis within the BNP paradigm addressing noise sources discussed above (items 1-4). In addition, we develop specialized computational schemes for both continuous and pulsed illumination for it to “cost less”.

Indeed, mitigating computational cost becomes critical especially with the added complexity of working within the BNP paradigm. This, in itself, warrants a detailed treatment of continuous and pulsed illumination analyses in two companion manuscripts.

To complement this theoretical framework, we also provide to the community a suite of programs called BNP-FRET written in the compiled language Julia for high performance. These freely available programs allow for comprehensive analysis of single photon smFRET time traces on immobilized molecules obtained with a wide variety of experimental setups.

In what follows, we will first present a forward model. Next, we will build an inverse strategy to learn full posteriors within the BNP paradigm. Finally, multiple examples are presented by applying the method to simulated data sets across different parameter regimes. Experimental data are treated in the two subsequent companion manuscripts.

## 2 Forward Model

### 2.1 Conventions

To be consistent throughout our three part manuscript, we precisely define some terms as follows

1. a macromolecular complex under study is always referred to as a *system*,
2. the configurations through which a system transitions are termed *system states*, typically labeled using *σ*,
3. FRET dyes undergo quantum mechanical transitions between *photophysical states*, typically labeled using *ψ*,
4. a system-FRET combination is always referred to as a *composite*,
5. a composite undergoes transitions among its *superstates*, typically labeled using *ϕ*,
6. all transition rates are typically labeled using λ,
7. the symbol *N* is generally used to represent the total number of discretized time windows, typically labeled with *n*, and
8. the symbol *w_n_* is generally used to represent the observations in the *n*-th time window.

### 2.2 smFRET Data

Here, we briefly describe the data collected from typical smFRET experiments analyzed by BNP-FRET. In such experiments, donor and acceptor dyes labeling a system can be excited using either continuous illumination, or pulsed illumination where short laser pulses arrive at regular time intervals. Moreover, acceptors can also be excited by nonradiative transfer of energy from an excited donor to a nearby acceptor. Upon relaxation, both donor and acceptor can emit photons collected by single photon detectors. These detectors record the set of photon arrival times and detection channels. We denote the arrival times by

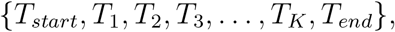

and detection channels with

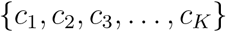

for a total number of *K* photons. In the equations above, *T_start_* and *T_end_* are experiment’s start and end times. Further, we emphasize here that the strategy used to index the detected photons above is independent of the illumination setup used.

Throughout the experiment, photon detection rates from the donor and acceptor dyes vary as the distance between them changes, due to the system kinetics. In cases where the distances form an approximately finite set, we treat the system as exploring a discrete system state space. The acquired FRET traces can then be analyzed to estimate the transition rates between these system states assuming a known model (*i.e.*, known number of system states). We will lift this assumption of knowing the model *a priori* in Sec. 3.2.

Cases where the system state space is continuous fall outside the scope of the current work and require extensions of Ref. [54] and Ref. [55] currently in progress.

In the following subsections, we present a physical model (forward model) describing the evolution of an immobilized system labeled with a FRET pair. We use this model to derive, step-by-step, the collected data’s likelihood given a choice of model parameters. Furthermore, given the mathematical nature of what is to follow, we will accompany major parts of our derivations with a pedagogical example of a molecule labeled with a FRET pair undergoing transitions between just two system states to demonstrate each new concept in example boxes.

### 2.3 Likelihood

To derive the likelihood, we begin by considering the stochastic evolution of an idealized system, transitioning through a discrete set of total *M_σ_* system states, {*σ*_1_,…, *σ_M_σ__*}, labeled with a FRET pair having *M_ψ_* discrete photophysical states, {*ψ*_1_,…, *ψ_M_ψ__*}, representing the fluorophores in their ground, excited, triplet, blinking, photobleached, or other quantum mechanical states. The combined system-FRET composite now undergoes transitions between *M_ϕ_* = *M_σ_* × *M_ψ_* superstates, {*ϕ*_1_,…, *φ_M_ϕ__*}, corresponding to all possible ordered pairs (*σ_j_,ψ_k_*) of the system and photophysical states. To be precise, we define *ϕ_i_* ≡ (*σ_j_, ψ_k_*), where *i* = (*j* – 1)*M_ψ_* + *k*.

Assuming Markovianity (memorylessness) of transitions among superstates, the probability of finding the composite in a specific superstate at a given instant evolves according to the master equation [40]

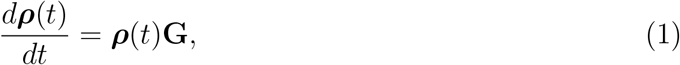

where the row vector ***ρ***(*t*) of length *M_ϕ_* has elements coinciding with probabilities for finding the system-FRET composite in a given superstate at time *t*. More explicitly, defining the photophysical portion of the probability vector ***ρ***(*t*) corresponding to system state *σ_i_* as

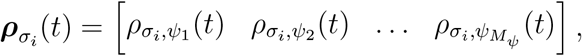

we can write ***ρ***(*t*) as

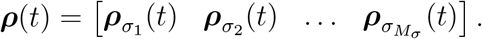

Furthermore, in the master equation above, **G** is the generator matrix of size *M_φ_* × *M_ϕ_* populated by all transition rates λ_*ϕ_i_*→*ϕ_j_*_ between superstates.

Each diagonal element of the generator matrix **G** corresponds to self-transitions and is equal to the negative sum of the remaining transition rates within the corresponding row. That is, λ*ϕ_i_*→*ϕ_i_* = -∑_*j*≠*i*_ λ_*ϕ_i_*→*ϕ_j_*_. This results in zero row-sums, assuring that ***ρ***(*t*) remains normalized at all times as described later in more detail (see Eq. 5). Furthermore, for simplicity, we assume no simultaneous transitions among system states and photophysical states as such events are rare (although the incorporation of these events in the model may be accommodated by expanding the superstate space). This assumption results in λ_(*ψ_i_,σ_j_*)→(*ψ_l_,σ_m_*)_ = 0 for simultaneous *i* ≠ *l* and *l* ≠ *m*, which allows us to simplify the notation further. That is, *λ*_(*ψ_i_,σ_j_*)→(*ψ_i_,σ_k_*)_ ≡ λ_*σ_j_*→*σ_k_*_ (for any *i*) and *λ*_(*ψ_i_,σ_j_*)→(*ψ_k_,σ_j_*)_ ≡ λ_*σ_j_,ψ_i_*→*ψ_k_*_ (for any *j*). This leads to the following form for the generator matrix containing blocks of exclusively photophysical and exclusively system transition rates, respectively

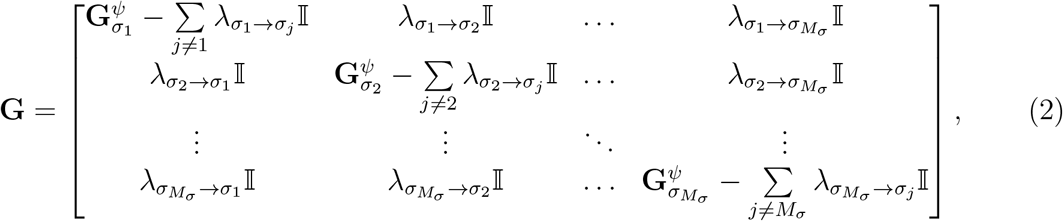

where the matrices on the diagonal 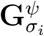 are the photophysical parts of the generator matrix for a system found in the *σ_i_* system state. Additionally, 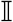 is the identity matrix of size *M_ψ_*.

For later convenience, we also organize the system transition rates λ_*σ_i_*→*σ_j_*_. in Eq. 2 as a matrix

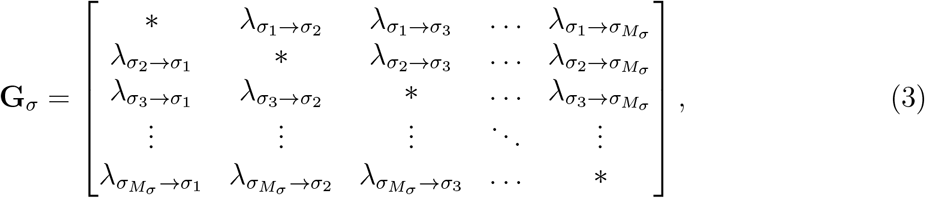

which we call system generator matrix.

Moreover, the explicit forms of 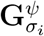 in Eq. 2 depend on the photophysical transitions allowed in the model. For instance, if the FRET pair is allowed to go from its ground photophysical state (*ψ*_1_) to the excited donor (*ψ*_2_) or excited acceptor (*ψ*_3_) states only, the matrix is given as

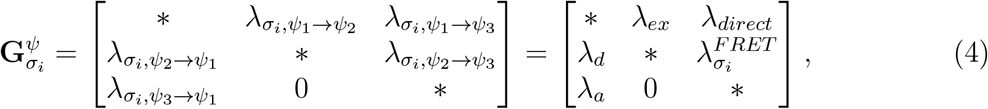

where the * along the diagonal represents the negative row-sum of the remaining elements, *λ_ex_* is the excitation rate, λ_*d*_ and λ_*α*_ are the donor and acceptor relaxation rates, respectively, and *λ_direct_* is direct excitation of the acceptor by a laser, and 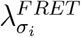 is the donor to acceptor FRET transition rate when the system is in its *i*-th system state. We note that only FRET transitions depend on the system states (identified by dye-dye separations) and correspond to FRET efficiencies given by

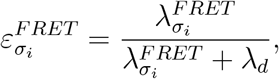

where the ratio on the right hand side represents the fraction of FRET transitions among all competing transitions out of an excited donor, that is, the fraction of emitted acceptor photons among total emitted photons.

With the generator matrix at hand, we now look for solutions to the master equation of Eq. 1. Due to its linearity, the master equation accommodates the following analytical solution:

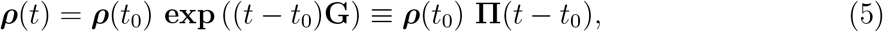

illustrating how the probability vector ***ρ***(*t*) arises from the propagation of the initial probability vector at time *t*_0_ by the exponential of the generator matrix (the propagator matrix **Π**(*t* – *t*_0_)). The exponential maps the transition rates λ*ϕ_i_*→*ϕ_j_* in the generator matrix to their corresponding transition probabilities Π_*ϕ_i_*→*ϕ_j_*_ populating the propagator matrix. The zero row-sums of the generator matrix guarantee that the resulting propagator matrix is stochastic (*i.e.*, has rows of probabilities that sum to unity, ∑_*j*_π_*ϕ_i_*→*ϕ_j_*_ = 1).

##### Example I: State Space and Generator Matrix

For a molecule undergoing transitions between its two conformations, we have *M_σ_* = 2 system states given as {*σ*_1_,*σ*_2_}. The photophysical states of the FRET pair labeling this molecule are defined according to whether the donor or acceptor are excited. Denoting the ground state by *G* and excited state by *E*, we can write all photophysical states of the FRET pair as {*ψ*_1_ = (*G, G*), *ψ*_2_ = (*E, G*), *ψ*_3_ = (*G,E*)}, where the first element in the ordered pair represents the donor state. Further, here, we assume no simultaneous excitation of the donor and acceptor owing to its rarity.

Next, we construct the superstate space with *M_ϕ_* = 6 ordered pairs {*ϕ*_1_ = (*ψ*_1_,*σ*_1_),*ϕ*_2_ = (*ψ*_2_,*σ*_1_), *ϕ*_3_ = (*ψ*_3_, *σ*_1_), *ϕ*_4_ = (*ψ*_1_, *σ*_2_), *ϕ*_5_ = (*ψ*_2_,*σ*_2_),*ϕ*_6_ = (*ψ*_3_,*σ*_2_)}. Finally, the full generator matrix for this setup reads

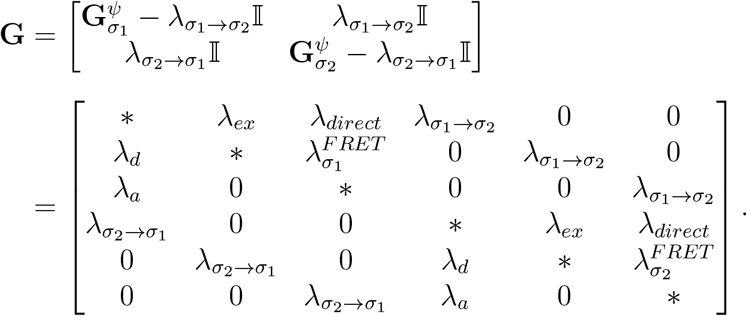

Both here, and in similar example boxes that follow, we choose values for rates commonly encountered in experiments [17]. We consider a laser exciting a donor at rate *λ_ex_* = 10 ms^-1^. Next, we suppose that the molecule switches between system states *σ*_1_ and *σ*_2_ at rates λ_*σ*_1_→*σ*_2__ = 2.0 ms^-1^ and λ_*σ*_2_→*σ*_1__ = 1 ms^-1^.

Furthermore, assuming typical lifetimes of 3.6 ns and 3.5 ns for the donor and acceptor dyes [17], their relaxation rates are, respectively, λ_*d*_ = 1/3.6 ns^-1^ and λ_*α*_ = 1/3.5 ns^-1^. We also assume that there is no direct excitation of the acceptor and thus λ_*direct*_ = 0. Next, we choose FRET efficiencies of 0.2 and 0.9 for the two system states resulting in 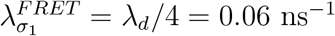 and 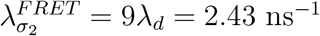.

Finally, these values lead to the following generator matrix (in ms^-1^ units)

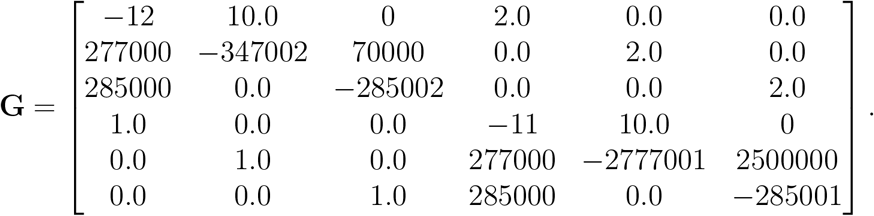

After describing the generator matrix and deriving the solution to the master equation, we continue by explaining how to incorporate observations into a likelihood.

In the absence of observations, any transition among the set of superstates are unconstrained. However, when monitoring the system using suitable detectors, observations rule out specific transitions at the observation time. For example, ignoring background for now, the detection of a photon from a FRET label identifies a transition from an excited photophysical state to a lower energy photophysical state of that label. On the other hand, no photon detected during a time period indicates the absence of radiative transitions or the failure of detectors to register such transition. Consequently, even periods without photon detections are informative in the presence of a detector. In other words, observations from a single photon smFRET experiment are continuous in that they are defined at every point in time.

Additionally, since smFRET traces report radiative transitions of the FRET labels at photon arrival times, uncertainty remains about the occurrences of unmonitored transitions (*e.g.*, between system states). Put differently, smFRET traces (observations) only partially specify superstates at any given time.

Now, to compute the likelihood for such smFRET traces, we must sum those probabilities over all trajectories across superstates (superstate trajectories) consistent with a given set of observations. Assuming the system ends in superstate *ϕ_i_* at *T_end_*, this sum over all possible trajectories can be very generally given by the element of the propagated vector ***ρ***(*T_end_*) corresponding to superstate *ϕ_i_*. Therefore, a general likelihood may be written as

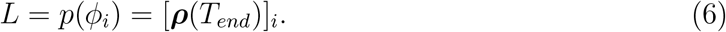

However, as the final superstate at time *T_end_* is usually unknown, we must therefore marginalize (sum) over the final superstate to obtain the following likelihood

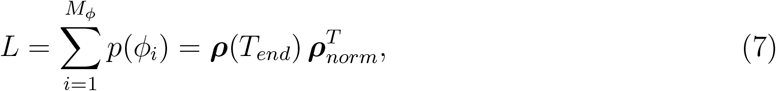

where all elements of the vector ***ρ***_*norm*_ are set to 1 as a means to sum the probabilities in vector ***ρ***(*T_end_*). In the following sections, we describe how to obtain concrete forms for these general likelihoods.

#### 2.3.1 Absence of observations

For pedagogical reasons, it is helpful to first look at the trivial case where a system-FRET composite evolves but no observations are made (due to a lack, say, of detection channels). In this case, all allowed superstate trajectories are possible between the start time of the experiment, *T_start_*, and end, *T_end_.* This is because the superstate cannot be specified or otherwise restricted at any given time by observations previously explained. Consequently, the probability vector ***ρ***(*t*) remains normalized throughout the experiment as no superstate trajectory is excluded. As such, the likelihood is given by summing over probabilities associated to the entire set of trajectories, that is,

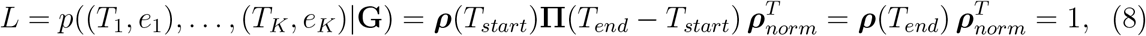

where {*e*_1_,…, *e_K_*} are the emission times of all emitted photons, not recorded due to lack of detection channels and thus not appearing on the right hand side of the expression.

In what follows, we describe how the probability vector ***ρ***(*t*) does not remain normalized as it evolves to ***ρ***(*T_end_*) when detectors partially collapse knowledge of the occupied superstate during the experiment. This results in a likelihood smaller than one. We do so for the conceptually simpler case of continuous illumination for now.

#### 2.3.2 Introducing observations

To compute the likelihood when single photon detectors are present, we start by defining a measurement model where the observation at a given time is dictated by ongoing transitions and detector features (*e.g.*, crosstalk, detector efficiency). As we will see in more detail later, if we describe the evolution of a system by defining its states at a discrete time points and these states are not directly observed, and thus hidden, then this measurement model adopts the form of a hidden Markov Model (HMM). Here, Markovianity arises when a given hidden state *only* depends on its immediate preceding hidden state. In such HMMs, an observation at a given time is directly derived from the concurrent hidden state.

As an example of an HMM, for binned smFRET traces, an observation is often approximated to depend only on the current hidden state. However, contrary to such a naive HMM, an observation in a single photon setup in a given time period depends on the current superstate and the immediate previous superstate. This naturally enforces a second order structure on the HMM where each observed random variable depends on two superstates as we demonstrate shortly. A similar HMM structure was noted previously to model a fluorophore’s photo switching behavior in Ref. [56].

Now, in order to address this observation model, we first divide the experiment’s time duration into *N* windows of equal size, *ϵ* = (*T_end_* – *T_start_*)/*N*. We will eventually take the continuum limit *ϵ* → 0 to recover the original system as described by the master equation. We also sum over all possible transitions between superstates within each window. These windows are marked by the times (see Fig. 2 (a))

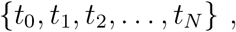

where the *n*-th window is given by (*t*_*n*–1_, *t_n_*) with *t*_0_ = *T_start_* and *t_N_* = *T_end_*. Corresponding to each time window, we have observations

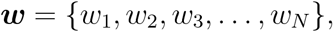

where *w_n_* = ⊘ if no photons are detected and 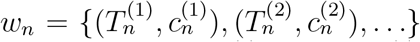 otherwise, with the *j*-th photon in a window being recorded by the channel 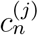 at time 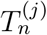. Note here that observations in a time window, being a continuous quantity, allow for multiple photon arrivals or none at all.

**Figure 2:**
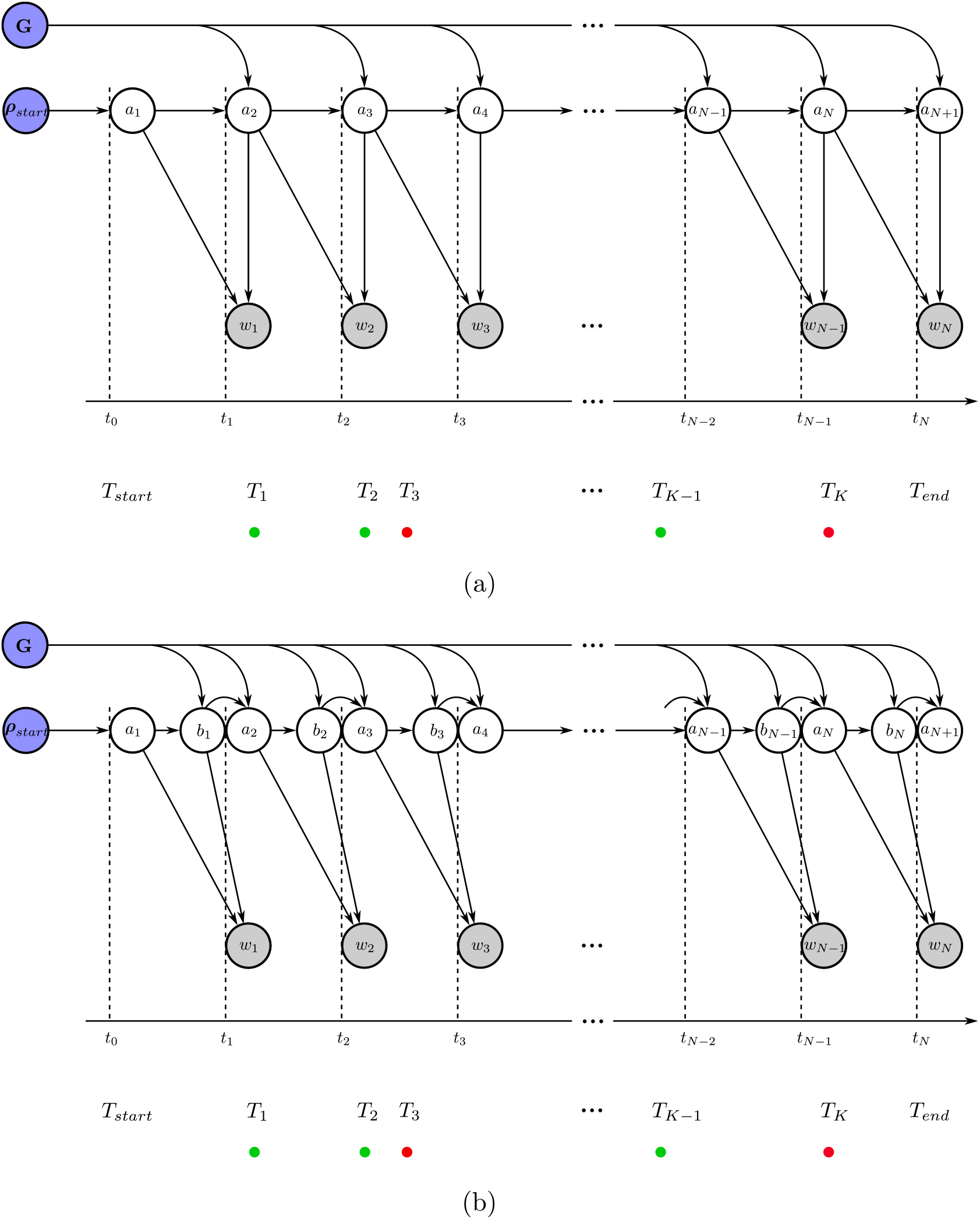
Graphical models depicting the random variables and parameters involved in the generation of photon arrival data for smFRET experiments. Circles shaded in blue represent parameters of interest we wish to deduce, namely transition rates and probabilities. The circles shaded in gray correspond to observations. The unshaded circles represent the superstates. The arrows reflect conditional dependence among these variables and colored dots represent photon arrivals. Going from panel (a) to (b), we convert the original HMM with a second order structure to a naive HMM where each observation only depends on one state.

As mentioned earlier, each of these observations originate from the evolution of the superstate. Therefore, we define superstates occupied at the beginning of each window as

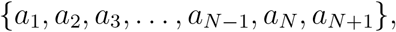

where *a_n_* is the superstate at the beginning of the *n*-th time window as shown in Fig. 2(a). The framework described here can be employed to compute the likelihood. However, the second order structure of the HMM leads to complications in these calculations. In the rest of this section, we first illustrate the mentioned complication using a simple example and then describe a solution to this issue.

###### Example II: Naive Likelihood Computation

Here, we calculate the likelihood for our two state system described earlier. For simplicity alone, we attempt the likelihood calculation for a time period spanning the first two time windows (*N* = 2) in Fig. 2(a). Within this period the system-FRET composite evolves from superstate *a*_1_ to *a*_3_ giving rise to observations *w*_1:2_. The likelihood for such a setup is typically obtained using a recursive strategy by marginalizing over superstates *a*_1:3_ (summing over all possible superstate trajectories)

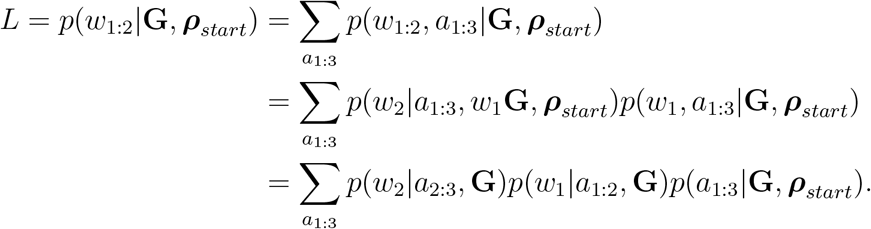

Here, we have applied the chain rule of probabilities in each step. Moreover, in the last step, we have only retained the parameters that are directly connected to the random variable on the left in each term, as shown by arrows in Fig. 2(a).

Now, for our two system state example, *a_n_* can be any of the six superstates *ϕ*_1:6_ (*M_ϕ_* = 6) given earlier. As such, the sum above contains 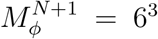 terms for such a simple example. For a large number of time windows, computing this sum becomes prohibitively expensive. Therefore, it is common to use a recursive approach to find the likelihood, only requiring 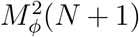 operations, as we will describe in the next section. However, due to our HMM’s second order structure, the two first terms (involving observations) in the above sum are conditioned on a mutual superstate *α*_2_ which forbids recursive calculations.

After describing the issue in computing the likelihood due to the second order structure of our HMM, we now describe a solution to this problem. As such, to simplify the likelihood calculation, we temporarily introduce superstates *b_n_* at the end of *n*-th window separated from superstate *a*_*n*+1_ at the beginning of (*n* + 1)-th window by a short time *τ* as shown in Fig. 2(b) during which no observations are recorded (inactive detectors). This procedure allows us to conveniently remove dependency of consecutive observations on a mutual superstate. That is, consecutive observations *w_n_* and *w*_*n*+1_ now do not depend on a common superstate *a*_*n*+1_, but rather on separated (*a_n_, b_n_*) pairs; see Fig. 2(b). The sequence of superstates now looks like (see Fig. 2b)

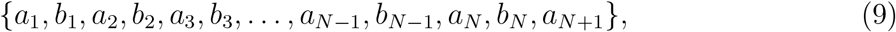

which now permits a recursive strategy for likelihood calculation as described in the next section. Furthermore, we will eventually take the *τ* → 0 limit to obtain the likelihood of the original HMM with the second order structure.

#### 2.3.3 Recursion formulas

We now have the means to compute the terminal probability vector ***ρ***_*end*_ = ***ρ***(*T_end_*) by evolving the initial vector ***ρ***_*start*_ = ***ρ***(*T_start_*). This is most conveniently achieved by recursively marginalizing (summing) over all superstates in Eq. 9 backwards in time, starting from the last superstate *a*_*N*+1_ as follows

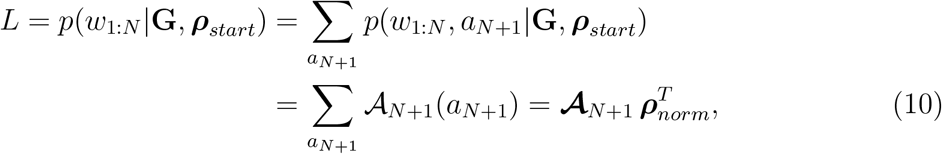

where 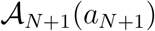 are elements of the vector 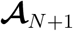 of length *M_ϕ_*, commonly known as a filter [57]. Moving backwards in time, the filter at the beginning of the *n*th time window, 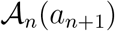, is related to the filter at the end of the *n*th window, 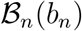, due to Markovianity, as follows

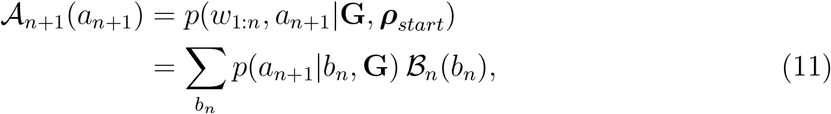

or in matrix notation as

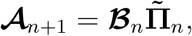

where *p*(*a*_*n*+1_|*b_n_*) are the elements of the transition probability matrix 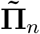 described in the next section. Again due to Markovianity, the filter at the end of the *n*th window, 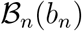, is related to the filter at the beginning of the same time window, 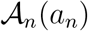, as

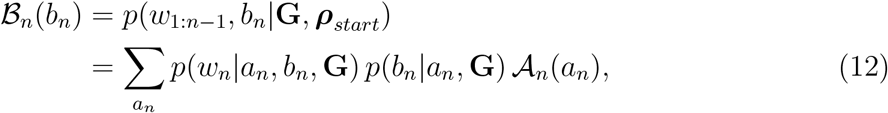

or in matrix notation as

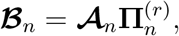

where the terms *p*(*w_n_*|*a_n_, b_n_*, **G**)*p*(*b_n_*|*a_n_*, **G**) populate the transition probability matrix 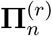 described in the next section. Here, we use the superscript (*r*) to denote that elements of this matrix include observation probabilities. We note here that the last filter in the recursion formula, 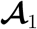, is equal to starting probability vector ***ρ***_*start*_ itself.

#### 2.3.4 Reduced propagators

To derive the different terms in the recursive filter formulas, we first note that the transition probabilities *p*(*a_n_*|*b*_*n*-1_, **G**) and *p*(*b_n_*|*a_n_*, **G**) do not involve observations. As such, we can use the full propagator as follows

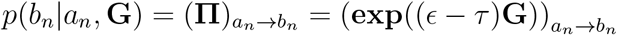

and

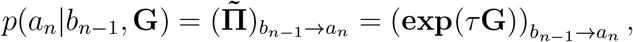

respectively. On the other hand, the term *p*(*w_n_*|*a_n_, b_n_*, **G**) includes observations which results in modification to the propagator by ruling out a subset of transitions. For instance, observation of a photon momentarily eliminates all nonradiative transitions. The modifications now required can be structured into a matrix **D**_*n*_ of the same size as the propagator with elements (**D**_*n*_)_*a*_*n*_→*b*_n__ = *p*(*w_n_*|*a_n_,b_n_*, **G**). We term all such matrices *detection* matrices. The product *p*(*w_n_*|*a_n_, b_n_*, **G**) *p*(*b_n_*|*a_n_*, **G**) in Eq. 12 can now be written as

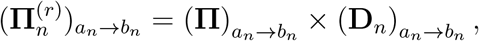

relating the modified propagator (termed reduced propagator and distinguished by the superscript (r) hereafter) 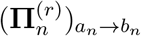 in the presence of observations to the full propagator (no observations). Plugging in the matrices introduced above into the recursive filter formulas (Eqs. 11-12), we obtain in matrix notation

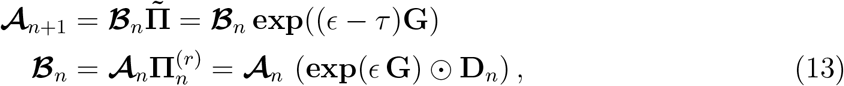

where the symbol ⊙ represents element-by-element product of matrices. Here, however, the detection matrices cannot yet be computed analytically as the observations *w_n_* allow for an arbitrary number of transitions within the finite time window (*t*_*n*-1_, *t_n_*). However, they become manageable in the limit that the time windows become vanishingly small, as we will demonstrate later.

#### 2.3.5 Likelihood for the HMM with second order structure

Now, inserting the matrix expressions for filters of Eq. 13 into the recursive formula likelihood Eq. 10, we arrive at

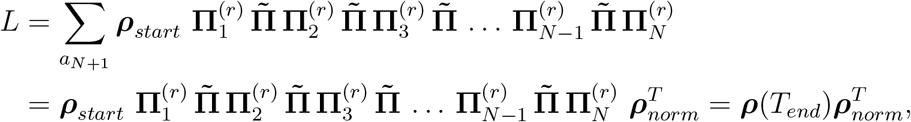

where, in the second step, we added a row vector of ones, ***ρ***_*norm*_ at the end to sum over all elements. Here, the superscript *T* denotes matrix transpose. As we now see, the structure of the likelihood above amounts to propagation of the initial probability vector ***ρ***_*start*_ to the final probability vector ***ρ***(*T_end_*) via multiple propagators corresponding to *N* time windows.

Now, under the limit *τ* → 0, we have

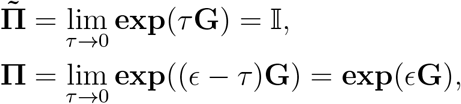

where 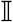 is the identity matrix. In this limit, we recover the likelihood for the HMM with a second order structure as

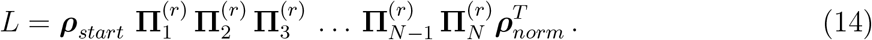

We note here that the final probability vector ***ρ***(*T_end_*) is not normalized to one upon propagation due to the presence of reduced propagators corresponding to observations. More precisely, the reduced propagators restrict the superstates evolution to only a subset of trajectories over a time window *ϵ* in agreement with the observation over this window. This, in turn, results in a probability vector whose elements sum to less than one. That is,

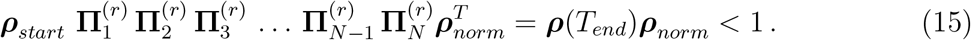

#### 2.3.6 Continuum limit

Up until now, the finite size of the time window *ϵ* allowed for an arbitrary number of transitions per time window (*t*_*n*-1_, *t_n_*), which hinders the computation of an exact form for the detection matrices. Here, we take the continuum limit, as the time windows become vanishingly small (that is, *ϵ* → 0 as *N* → ∞). Thus, no more than one transition is permitted per window. This allows us to fully specify the detection matrices **D**_*n*_.

To derive the detection matrices, we first assume ideal detectors with 100% efficiency and include detector effects in the subsequent sections (see Sec. 2.4). In such cases, the absence of photon detections during a time window, while detectors are active, indicates that only nonradiative transitions took place. Thus, only nonradiative transitions have nonzero probabilities in the detection matrices. As such, for evolution from superstate *a_n_* to *b_n_*, the elements of the nonradiative detection matrix, **D**^*non*^, are given by

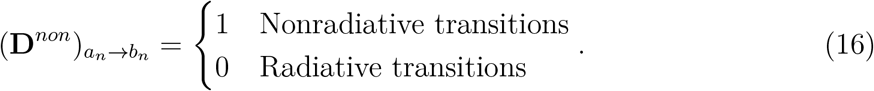

On the other hand, when the *k*-th photon is recorded in a time window, only elements corresponding to radiative transitions are nonzero in the detection matrix denoted by 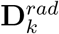 as

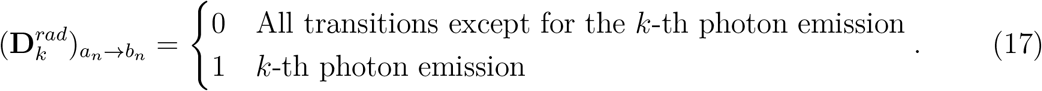

Here, we note that the radiative detection matrices have zeros along their diagonals, since self-transitions are nonradiative.

We can now define the reduced propagators corresponding to the nonradiative and radiative detection matrices, **D**^*non*^ and 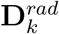, using the Taylor approximation 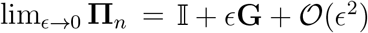 as

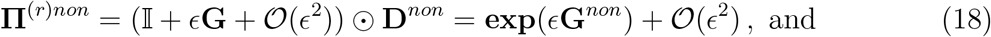

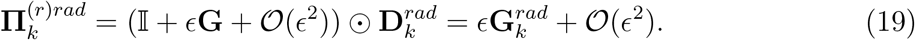

In the equations above, **G**^*non*^ = **G** ⊙ **D**^*non*^ and 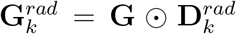, where the symbol ⊙ represents an element-by-element product of the matrices. Furthermore, the product between the identity matrix and 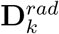 above vanishes in the radiative propagator due to zeros along the diagonals of 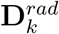.

###### Example III: Detection Matrices

For our example with two system states described earlier, the detection matrices of Eqs. 16-17 take simple forms. The radiative detection matrix has the same size as the generator matrix with nonzero elements wherever there is a rate associated to a radiative transition

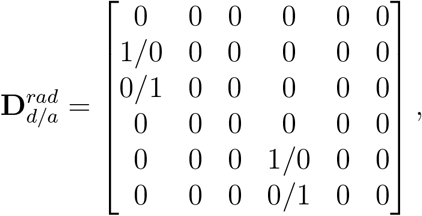

where the subscripts *d* and *a*, respectively, denote photon detection in donor and acceptor channels. Similarly, the nonradiative detection matrix is obtained by setting all elements of the generator matrix related to radiative transitions to zero and the remaining to one as

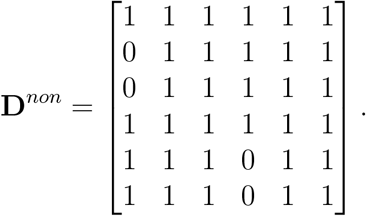

#### 2.3.7 Final likelihood

With the asymptotic forms of the reduced propagators in Eq. 14 now defined in the last subsection, we have all the ingredients needed to arrive at the final form of the likelihood.

To do so, we begin by considering the period right after the detection of the (*k* – 1)th photon until the detection of the *k*th photon. For this time period, the nonradiative propagators in Eq. 14 can now be easily merged into a single propagator 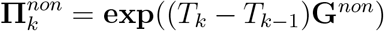, as the commutative arguments of the exponentials can be readily added. Furthermore, at the end of this interphoton period, the radiative propagator 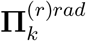 marks the arrival of the *k*-th photon. The product of these two propagators

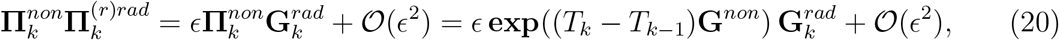

now governs the stochastic evolution of the system-FRET composite during that interphoton period.

Inserting Eq.20 for each interphoton period into the likelihood for the HMM with second order structure in Eq. 14, we finally arrive at our desired likelihood

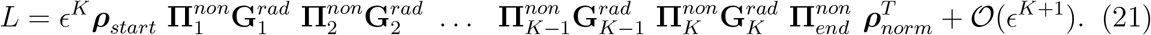

This likelihood has the same structure as shown by Gopich and Szabo in Ref. [40].

###### Example IV: Propagator and Likelihood

Here, we consider a simple FRET trace where two photons are detected at times 0.05 ms and 0.15 ms in the donor and acceptor channels, respectively. To demonstrate the ideas developed so far, we calculate the likelihood of these observations as (see Eq. 21)

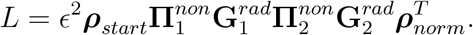

To do so, we first need to calculate 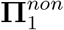 using the nonradiative detection (**D**^*non*^) and generator (**G**) matrices found in the previous example boxes

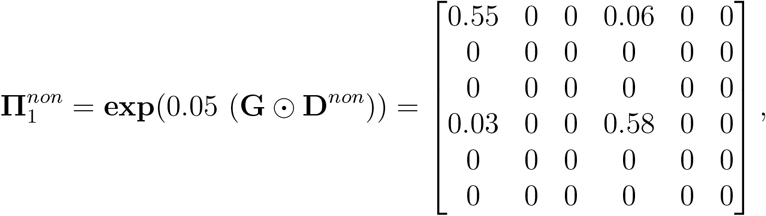

and similarly

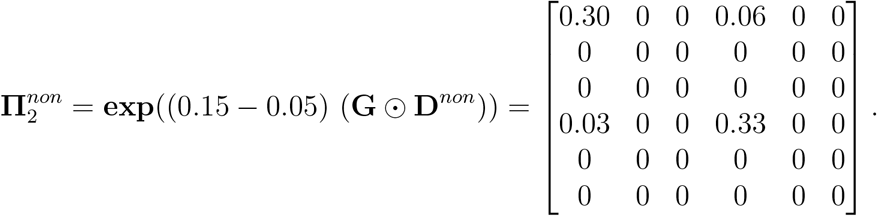

Next, we proceed to calculate 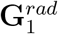 and 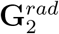. Remembering that the first photon was detected in the donor channel, we have (in ms^-1^ units)

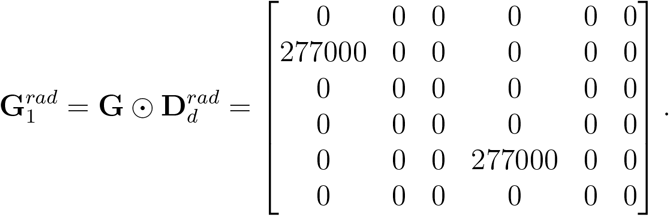

Similarly, since the second photon was detected in the acceptor channel, we can write (in ms^-1^ units)

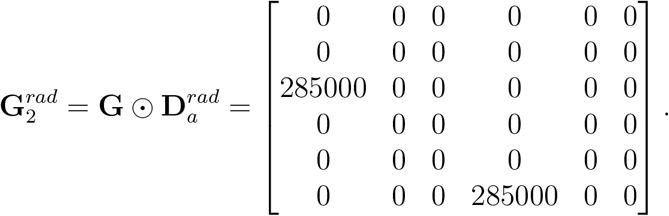

We also assume that the system is initially in the superstate *ϕ*_1_ giving ***ρ***_*start*_ = [1, 0, 0, 0, 0, 0]. Finally, putting everything together, we can find the likelihood as *L* = 3.06*ϵ*^2^ where e is a constant and does not contribute to parameter estimations as we will show later.

#### 2.3.8 Effect of binning single photon smFRET data

When considering binned FRET data, the time period of an experiment (*T_end_* – *T_start_*) is typically divided into a finite number (= *N*) of equally sized (= *ϵ*) time windows (bins), and the photon counts (intensities) in each bin are recorded in the detection channels. This is in constrast to single photon analysis where individual photon arrival times are recorded. To arrive at the likelihood for such binned data, we start with the single photon likelihood derived in Eq. 15 where *ϵ* is not infinitesimally small, that is,

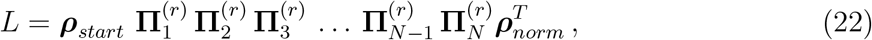

where

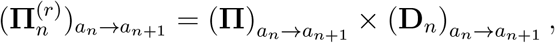

or in the matrix notation

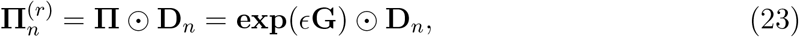

where **D**_*n*_ is the detection matrix introduced in Sec. 2.3.4.

Next, we must sum over all superstate trajectories that may give rise to the recorded photon counts (observations) in each bin. However, such a sum is challenging to compute analytically and has been attempted in [38]. Here, we will only show likelihood computation under commonly applied approximations/assumptions when analyzing binned smFRET data, which are: 1) bin size *ϵ* is much smaller than typical times spent in a system state or, in other words, for a system transition rate λ_*σ_i_*→*σ_j_*_, we have *ϵ*λ_*σ_i_*→*σ_j_*_ ≪ 1; and 2) excitation rate *λ_ex_* is much slower than dye relaxation and FRET rates, or in other words, interphoton periods are much larger than the excited state lifetimes.

The first assumption is based on realistic situations where system kinetics (at seconds timescale) are many orders of magnitude slower than the photophysical transitions (at nanoseconds timescale). This timescale separation allows us to simplify the propagator calculation in Eq. 23. To see that, we first separate the system transition rates from photophysical transition rates in the generator matrix as

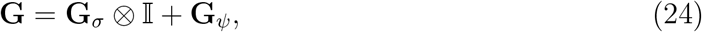

where ⊗ denotes a tensor product, **G**_*σ*_ is the portion of generator matrix **G** containing only system transition rates previously defined in Eq. 3, and **G**_*ψ*_ is the portion containing only photophysical transition rates, that is,

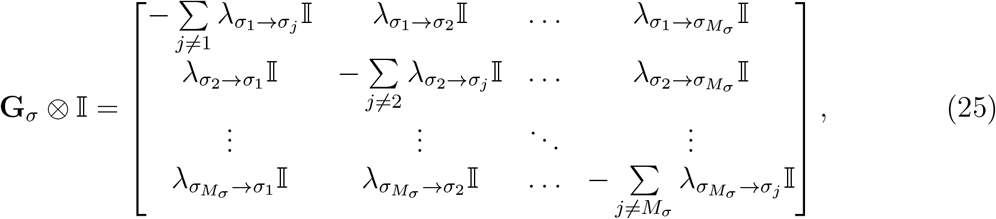

and

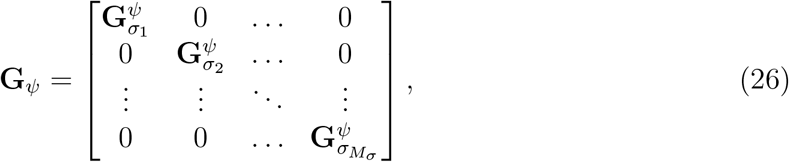

where 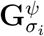 is the photophysical generator matrix corresponding to system state *σi* given in Eq. 4.

Now plugging Eq. 24 into the full propagator **Π** = **exp**(*ϵ***G**) and applying the famous Zassenhaus formula for matrix exponentials, we get

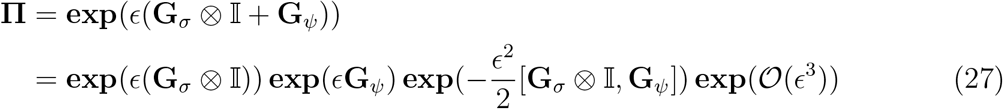

where the square brackets represent the commutator of the constituting matrices and the last term represents the remaining exponentials involving higher order commutators. Furthermore, the commutator 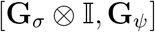 results in a very sparse matrix given by

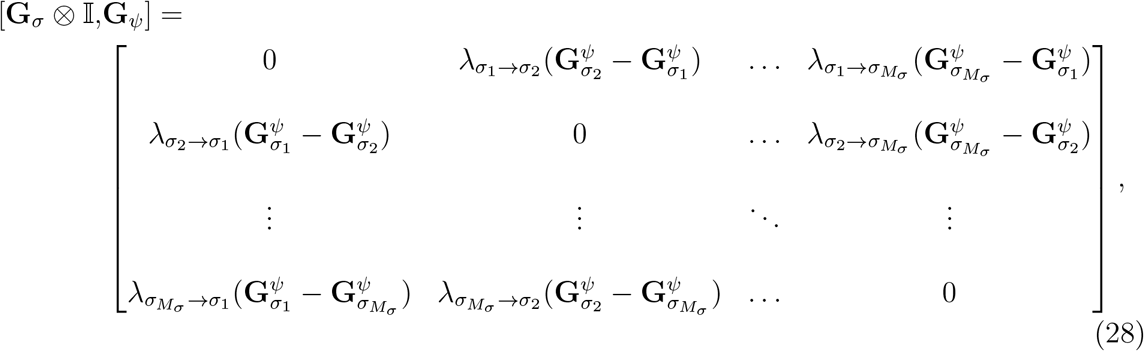

where

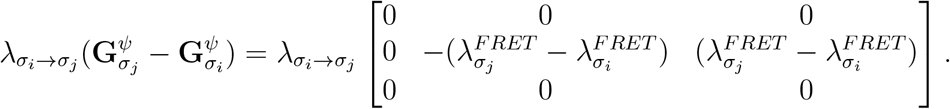

Now, the propagator calculation in Eq. 27 simplifies if the commutator 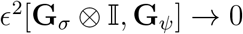, implying that either the bin size *ϵ* is very small such that *ϵ*λ_*σ_i_*→*σ_j_*_ ≪ 1 (our first assumption) or FRET rates/efficiencies are almost indistinguishable 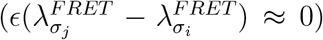. Under such conditions, the system state can be assumed to stay constant during a bin, with system transitions only occurring at the ends of bin periods. Furthermore, the full propagator **Π** in Eq. 27 can now be approximated as

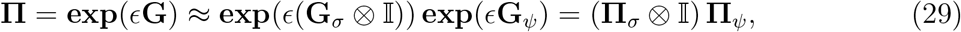

where the last equality follows from the block diagonal form of 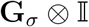 given in Eq. 25 and **Π**_*σ*_ = **exp**(*ϵ***G**_*σ*_) is the system transition probability matrix (propagator) given as

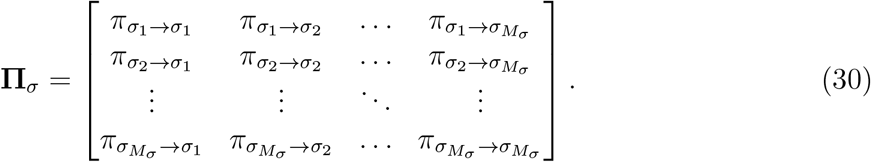

Moreover, **Π**_*ψ*_ = **exp**(*ϵ***G**_*ψ*_) is the photophysical transition probability matrix (propagator) as

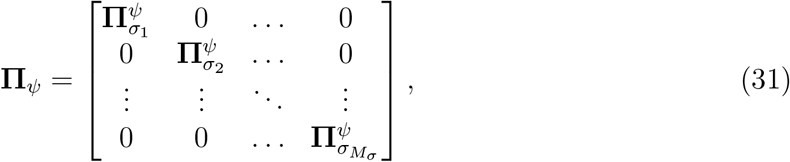

where the elements are given as 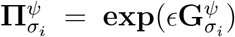. Furthermore, because of the block diagonal structure of **Π**_*ψ*_, the matrix multiplication in Eq. 29 results in

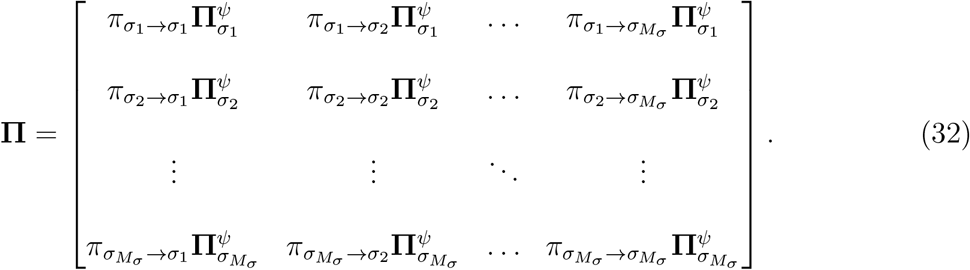

After deriving the full propagator **Π** for the time period *ϵ* (bin) under our first assumption, we now proceed to incorporate observations during this period via detection matrices **D**_*n*_ to compute the reduced propagator of Eq. 23. To do so, we now apply our second assumption of relatively slower excitation rate λ_*ex*_. This assumption implies that interphoton periods are dominated by the time spent in the ground state of the FRET pair and are distributed according to a single exponential distribution, **Exponential**(λ_*ex*_). Consequently, the total photon counts per bin follow a Poisson distribution, **Poisson**(eλ_*ex*_), independent of the photophysical portion of the photophysical trajectory taken from superstate *a_n_* to *a*_*n*+1_.

Now, the first and the second assumptions imply that the observation during the *n*-th bin only depends on the system state *s_n_* (or the associated FRET rate 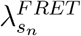). As such we can approximate the detection matrix elements as

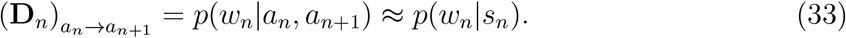

Using these approximations, the reduced propagator in Eq. 23 can now be written as

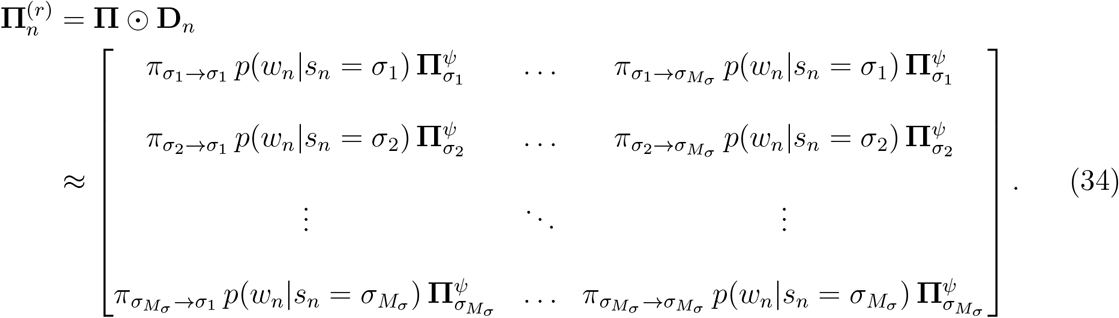

Next, to compute the likelihood for the *n*-th bin, we need to sum over all possible superstate trajectories within this bin as

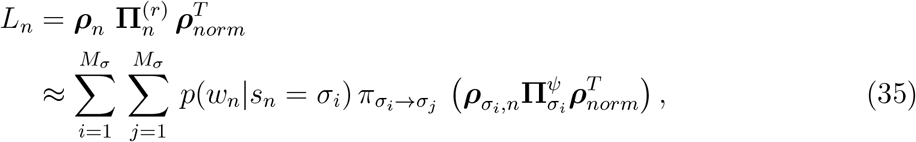

where ***ρ***_*n*_ is a normalized row vector populated by probabilities of finding the system-FRET composite in the possible superstates at the beginning of the *n*-th bin. Furthermore, we have written portions of ***ρ***_*n*_ corresponding to system state *σ_i_* as ***ρ***_*σ_i_*, *n*_. To be more explicit, we have

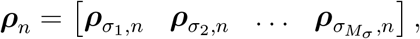

following the convention in Sec. 2.3 and using *n* to now represent time *t_n_*.

Moreover, since each row of 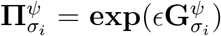 sums to one, we have 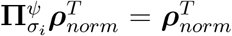, which simplifies the bin likelihood of Eq. 35 to

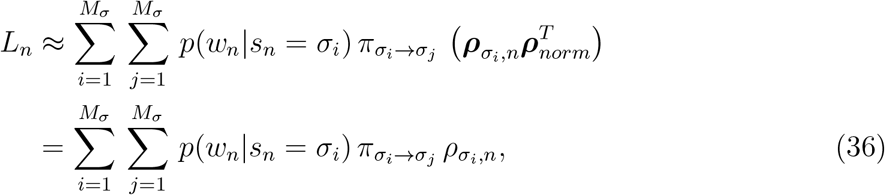

where we have defined 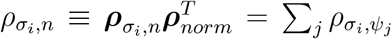 as the probability of the system to occupy system state *σ_i_*. We can also write the previous equation in the matrix form as

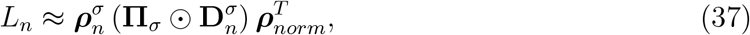

where 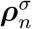 is a row vector of length *M_σ_* (number of system states) populated by *ρ_σ_i,n__* for each system state, and 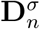, in the same spirit as **D**_*n*_, is a detection matrix of dimensions *M_σ_* × *M_σ_* populated by observation probability *p*(*w_n_*|*s_n_* = *σ_i_*) in each row corresponding to system state *σ_i_*. Furthermore, defining 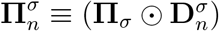, we note here that 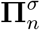 propagates probabilities during the *n*-th bin in a similar manner as the reduced propagators 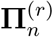 of Eq. 23.

Therefore, we can now multiply these new propagators for each bin to approximate the likelihood of Eq. 22 as

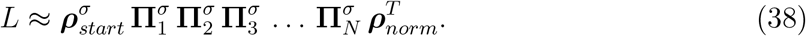

where 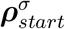 is a row vector, similar to 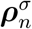, populated by probabilities of being in a given system state at the beginning of an experiment.

To conclude, our two assumptions regarding system kinetics and excitation rate allow us to significantly reduce the dimensions of the propagators. This, in turn, leads to much lowered expense for likelihood computation. However, cheaper computation comes at the expense of requiring large number of photon detections or excitation rate per bin to accurately determine FRET efficiencies (identify system states) since we marginalize over photophysics in each bin. Such high excitation rates lead to faster photobleaching and increased phototoxicity, and thereby much shorter experiment durations. As we will see in Sec. 2.5.1, this problem can be mitigated by using pulsed illumination, where the likelihood takes a similar form as Eq. 38, but FRET efficiencies can be accurately estimated from the measured microtimes.

### 2.4 Detection Effects

In the previous section, we assumed idealized detectors to illustrate basic ideas on detection matrices. However, realistic FRET experiments must typically account for detector nonidealities. For instance, many emitted photons may simply go undetected when the detection efficiency of single photon detectors, *i.e.*, the probability of an incident photon being successfully registered, is less than one due to inherent nonlinearities associated with the electronics [22] or the use of filters in cases of polarized fluorescent emission [58, 59]. Additionally, donor photons may be detected in the channel reserved for acceptor photons or vice-versa due to emission spectrum overlap [60]. This phenomenon, commonly known as crosstalk, crossover, or bleedthrough, can significantly affect the determination of quantities such as transition rates and FRET efficiencies as we demonstrate later in our results. Other effects adding noise to fluorescence signal include dark current (false signal in the absence of incident photons), dead time (the time a detector takes to relax back into its active mode after a photon detection), and timing jitter or IRF (stochastic delay in the output signal after a detector receives a photon) [22]. In this section, we describe the incorporation of all such effects into our model except dark current and background emissions which require more careful treatment and will be discussed in Sec. 2.6.

#### 2.4.1 Crosstalk and detection efficiency

Noise sources such as crosstalk and detection efficiency necessarily result in photon detection being treated as a stochastic process. Both crosstalk and detection efficiency can be included into the propagators in both cases by substituting the zeros and ones, appearing in the ideal radiative and nonradiative detection matrices (Eqs. 16-17), with probabilities between zero and one. In such a way, the resulting propagators obtained from these detection matrices, in turn, incorporate into the likelihood the effects of crosstalk and detection efficiency into the model.

Here, in the presence of crosstalk, for clarity, we add a superscript to the radiative detection matrix of Eq. 17 for the *k*-th photon, 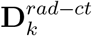. The elements of this detection matrix for the *a_n_* → *b_n_* transition, when a photon intended for channel *j* is registered in channel *i* reads

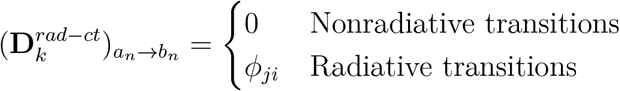

where *ϕ_ji_* is the probability for this event (upon transition from superstate *a_n_* to *b_n_*). Further, detector efficiencies can also be accounted for in these probabilities in order to represent the combined effects of crosstalk, arising from spectral overlap, and absence of detection channel registration. When we do so, we recover ∑_*i*_ *ϕ_ji_* ≤ 1 (for cases where *i* and *j* can be both the same or different), as not all emitted photons can be accounted for by the detection channels.

This new detection matrix above results in the following modification to the radiative propagator of Eq. 19 for the *k*-th photon

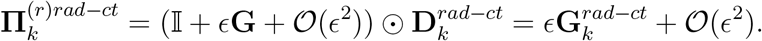

The second equality above follows by recognizing that the identity matrix multiplied, elementwise, by 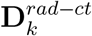 is zero. By definition, 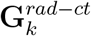 is the remaining nonzero product.

On the other hand, for time periods when no photons are detected, the nonradiative detection matrices in Eq. 16 become

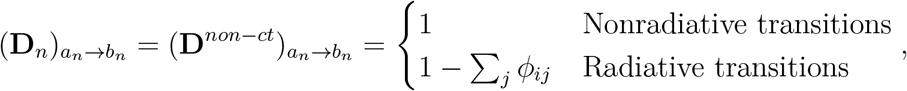

where the sum gives the probability of the photon intended for channel *i* to be registered in any channel. The nonradiative propagator of Eq. 18 for an infinitesimal period of size *ϵ* in the presence of crosstalk and inefficient detectors is now

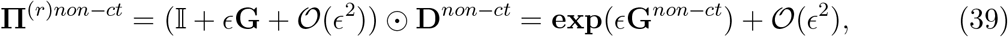

where **G**^*non-ct*^ = **G** ⊙ **D**^*non-ct*^. With the propagators incorporating crosstalk and detection efficiency now defined, the evolution during an interphoton period between the (*k* – 1)-th photon and the k-th photon of size (*T_k_* – *T*_*k*-1_) is now governed by the product

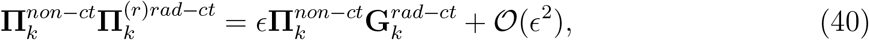

where the nonradiative propagators in Eq. 39 have now been merged into a single propagator 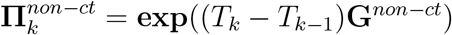 following the same procedure as Eq. 20.

Finally, inserting Eq. 40 for each interphoton period into the likelihood of Eq. 14, we arrive at the final likelihood incorporating crosstalk and detection efficiency as

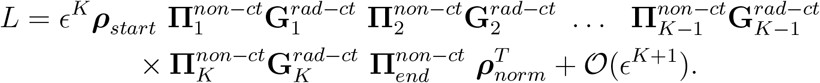

After incorporating crosstalk and detector efficiencies into our model, we briefly explain the calibration of the crosstalk probabilities/detection efficiencies *ϕ_ij_*. To calibrate these parameters, two samples, one containing only donor dyes and another containing only acceptor dyes, are individually excited with a laser under the same power to determine the number of donor photons 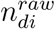 and number of acceptor photons 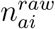 detected in channel i.

From photon counts recorded for the donor only sample, assuming ideal detectores with 100% efficiency, we can compute the crosstalk probabilities for donor photons going to channel *i, ϕ_di_*, using the photon count ratios as 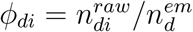 where 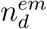 is the absolute number of emitted donor photons. Similarly, crosstalk probabilities for acceptor photons going to channel *i, ϕ_ai_*, can be estimated as 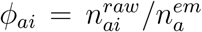 where 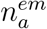 is the absolute number of emitted acceptor photons. In the matrix form, these crosstalk factors for a two-detector setup can be written as

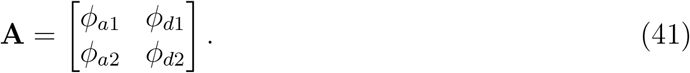

Using this matrix, for the donor only sample, we can now write

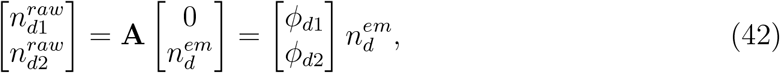

and similarly for the acceptor only sample

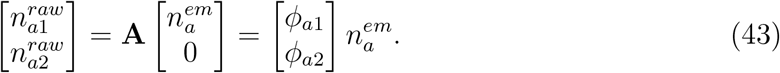

However, it is difficult to estimate the absolute number of emitted photons 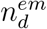 and 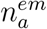 experimentally, and therefore the crosstalk factors in **A** can only be determined up to multiplicative factors of 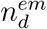 and 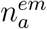.

Since scaling the photon counts in an smFRET trace by an overall constant does not affect the FRET efficiency estimates (determined by photon count ratios) and escape rates (determined by changes in FRET efficiency), we only require crosstalk factors up to a constant as in the last equation.

For this reason, one possible solution toward determining the matrix elements of **A** up to one multiplicative constant is to first tune dye concentrations such that the ratio 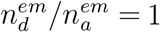, which can be accomplished experimentally. This allows us to write the crosstalk factors in the matrix form up to a constant as follows

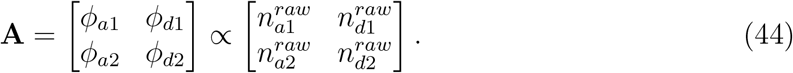

It is common to set the multiplicative factor in Eq. 44 by the total donor photons counts 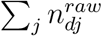 to give

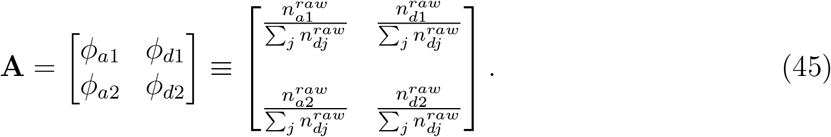

We note that from the convention adopted here, we have *ϕ*_*d*1_ + *ϕ*_*d*2_ = 1.

Furthermore, in situations where realistic detectors affect the raw counts, the matrix elements of **A** as computed above automatically incorporate the effects of detector inefficiencies including the fact that ∑_*j*_ *ϕ_j_* ≤ 1.

Additionally, the matrix **A** can be further generalized to account for more than two detectors by appropriately expanding the size of the matrix dimensions to coincide with the number of detectors. Calibration of the matrix elements then follows the same procedure as above.

Now, in performing single photon FRET analysis, we will use directly the elements of **A** in constructing our measurement matrix. However, it is also common, to compute the matrix elements of **A** from what is termed the route correction matrix (RCM) [61] typically used in binned photon analysis. The RCM is defined as the inverse of **A** to obtain corrected counts 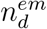 and 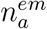 up to a proportionality constant as

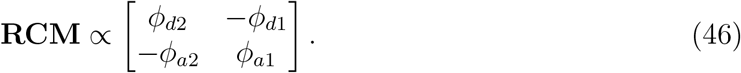

###### Example V: Detection Matrices with Crosstalk and Detector Efficiencies

For our example with two system states, we had earlier shown detection matrices for ideal detectors. Here, we incorporate crosstalk and detector efficiencies into these matrices. Moreover, we assume a realistic RCM [62] given as

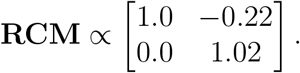

However, following the convention of Eq. 45, we scale the matrix provided by a sum of absolute values of its first row elements, namely 1.22, leading to effective crosstalk factors *ϕ_ij_* given as

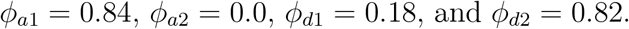

As such, these values imply approximately 18% crosstalk from donor to acceptor channel and 84% detection efficiency for acceptor channel without any crosstalk using the convention adopted in Eq. 45. Now, we modify the ideal radiative detection matrices by replacing their nonzero elements with the calibrated *ϕ_ij_*’s above

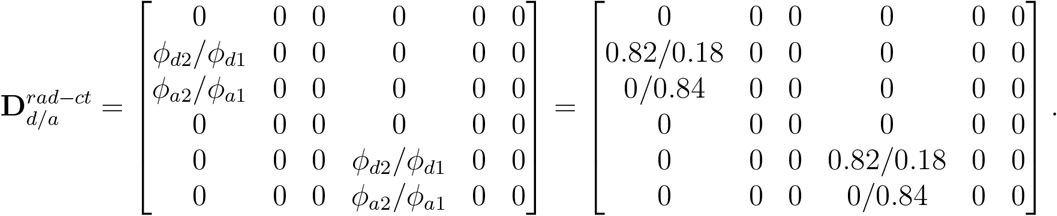

Similarly, we modify the ideal nonradiative detection matrix by replacing the zero elements by 1 – *ϕ*_*a*1_ – *ϕ*_*a*2_ = 0.16 and 1 – *ϕ*_*d*2_ – *ϕ*_*d*1_ = 0 as follows

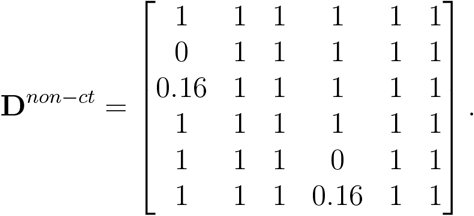

#### 2.4.2 Effects of detector dead time

Typically, a detection channel *i* becomes inactive (dead) after the detection of a photon for a period *δ_i_* as specified by the manufacturer. Consequently, radiative transitions associated with that channel cannot be monitored during that period.

To incorporate this detector dead period into our likelihood model, we break an interphoton period between the (*k* – 1)-th and *k*-th photon into two intervals: the first interval with an inactive detector and the second one when the detector is active. Assuming that the (*k* – 1)-th photon is detected in the *i*th channel, the first interval is thus *δ_ik_* long. As such, we can define the detection matrix for this interval as

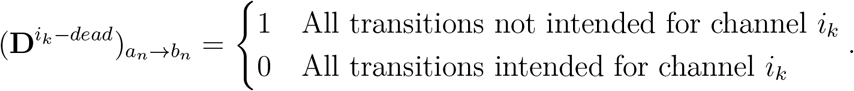

Next, corresponding to this detection matrix, we have the propagator

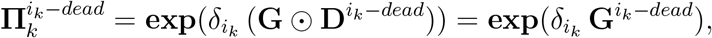

that evolves the superstate during the detector dead time. This propagator can now be used to incorporate the detector dead time into Eq. 20 to represent the evolution during the period between the (*k* – 1)-th and *k*-th photons as

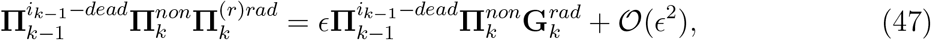

where 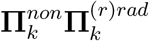 describes the evolution when the detector is active.

Finally, inserting Eq. 47 for each interphoton period into the likelihood for the HMM with a second order structure in Eq. 14, we arrive at the following likelihood that includes detector dead time

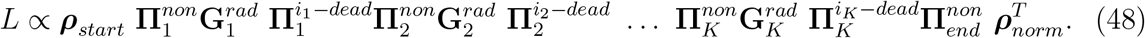

To provide an explicit example on the effect of the detector dead time on the likelihood, we take a detour for pedagogical reasons. In this context, we consider a very simple case of one detection channel (dead time *δ*) observing a fluorophore with two photophysical states, ground (*ψ*_1_) and excited (*ψ*_2_), illuminated by a laser. The data in this case contains only photon arrival times

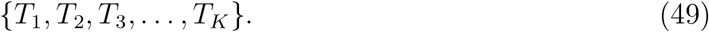

The generator matrix containing the photophysical transition rates for this setup is

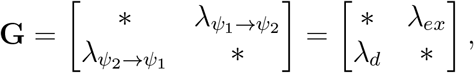

where the * along the diagonal represents the negative row-sum of the remaining elements, λ_*ex*_ is the excitation rate, and λ_*d*_ is the donor relaxation rate.

Here, all transitions are possible during detector dead times as there are no observations. As such, the dead time propagators in the likelihood (Eq. 48) are simply expressed as exponentials of the full generator matrix, that is, 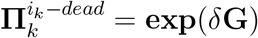, leaving the normalization of the propagated probability vector ***ρ*** unchanged, *e.g.*, just as we had seen in Eq. 8.

As we will see, these dead times, similar to detector inefficiencies, simply increase our uncertainty over parameters we wish to learn, such as kinetics, by virtue of providing less information. By contrast, background emissions and cross talk, provide false information. However, the net effect is the same: all noise sources increasing uncertainty.

#### 2.4.3 Adding the detection IRF

Due to various sources of noise impacting the detection timing electronics (also known as jitter), the time elapsed between photon arrival and detection is itself a hidden (latent) random variable [22]. Under continuous illumination, we say that this stochastic delay in time is sampled from a probability density, *f*(*τ*), termed the *detection* instrument response function (IRF). To incorporate the detection IRF into the likelihood of Eq. 48, we convolute the propagators with *f*(*τ*) as follows

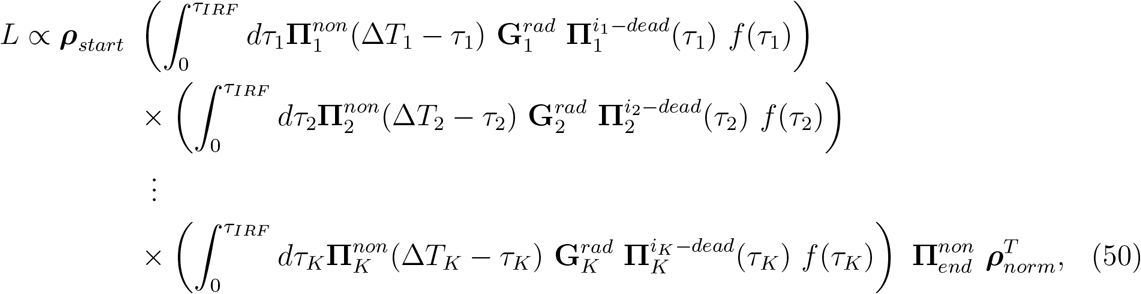

where we have used dead time propagators 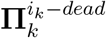 to incorporate detector inactivity during the period between photon reception and detector reactivation. Moreover, we have 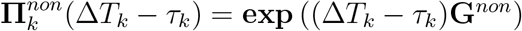 as described in Eq. 18.

To facilitate the computation of this likelihood, we use the fact that typical acquisition devices record at discrete (but very small) time intervals. For instance, a setup with the smallest acquisition time of 16 ps and a detection IRF distribution that is approximately 100 ps wide will have the detection IRF spread over, roughly, six acquisition periods. This allows each convolution integral to be discretized over the six acquisition intervals and computed in parallel, thereby avoiding extra real computational time associated to this convolution other than the overhead associated with parallelization.

### 2.5 Illumination Features

After discussing detector effects, we continue here by further considering different illumination features. For simplicity alone, our likelihood computation until now assumed continuous illumination with a uniform intensity. More precisely, the element λ_*ex*_ of the generator matrix in Eq. 4 was assumed to be time-independent. Here, we generalize our formulation and show how other illumination setups (such as pulsed illumination and alternating laser excitation, ALEX [63]) can be incorporated into the likelihood by simply assigning a time dependence to the excitation rate λ_*ex*_(*t*).

#### 2.5.1 Pulsed illumination

Here, we consider an smFRET experiment where the FRET pair is illuminated using a laser for a very short period of time (a pulse), *δ_pulse_*, at regular intervals of size *τ*; see Fig. 3(a). Now, as in the case of continuous illumination with constant intensity, the likelihood for a set of observations acquired using pulsed illumination takes a similar form to Eq. 21 involving products of matrices as follows

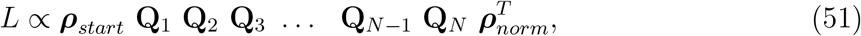

where **Q**_*n*_, with *n* = 1,…, *N*, denotes the propagator evolving the superstate during the *n*-th interpulse period between the (*n* – 1)-th and the *n*-th pulse.

**Figure 3:**
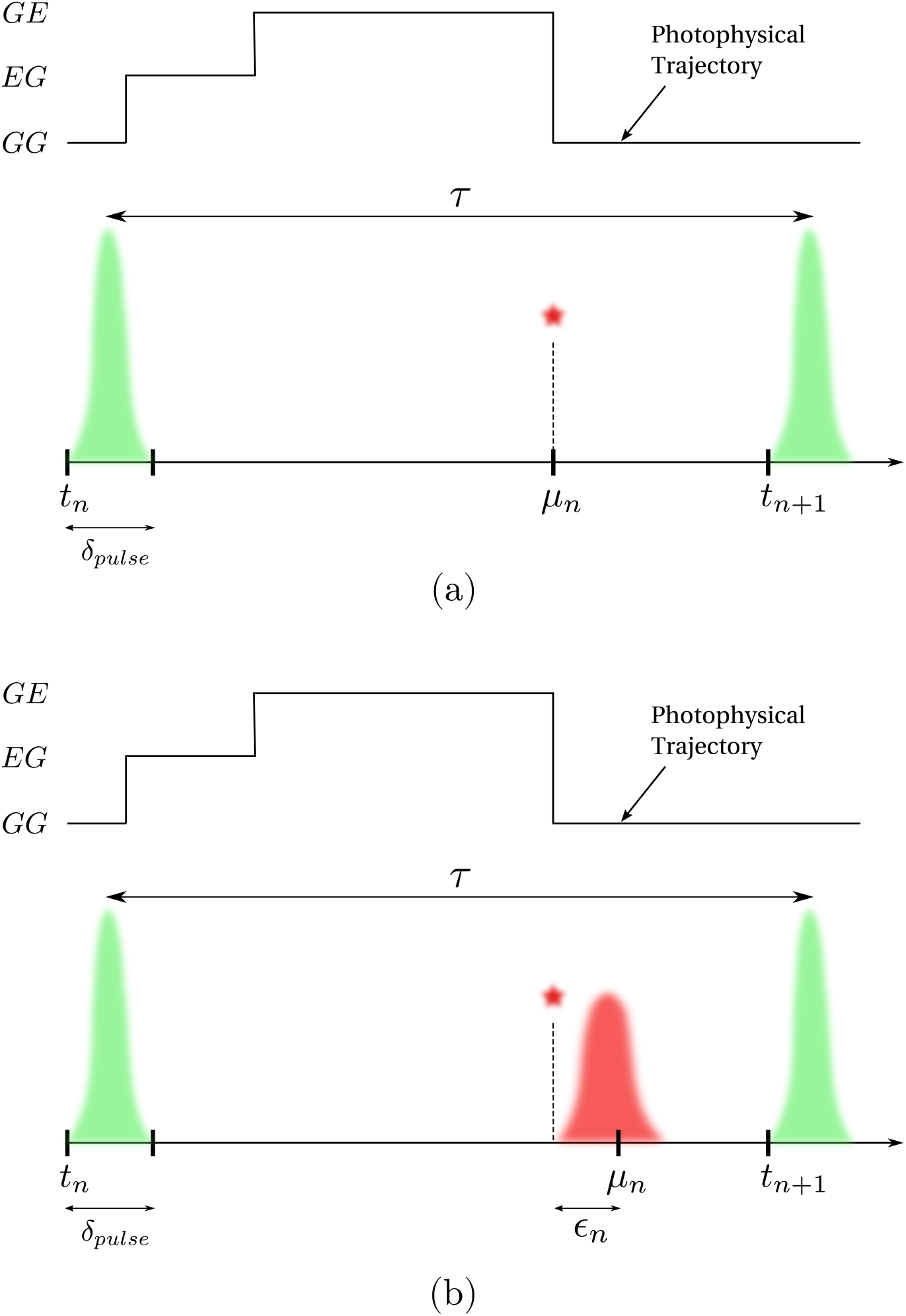
Events over a pulsed illumination experiment pulse window. Here, the beginning of the *n*-th interpulse window of size *τ* is marked by time *t_n_*. The FRET labels originally in the state GG (donor and acceptor, respectively, in ground states) are excited by a high intensity burst (shown by the green) to the state EG (only donor excited) for a very short time *δ_pulse_*. If FRET occurs, the donor transfers its energy to the acceptor and resides in the ground state leaving the FRET labels in the GE state (only acceptor excited). The acceptor then emits a photon to be registered by the detector at microtime *μ_n_*. When using ideal detectors, the microtime is the same as the photon emission time as shown in panel (a). However, when the timing hardware has jitter (shown in red), a small delay *ϵ_n_* is added to the microtime as shown in panel (b).

To derive the structure of **Q**_*n*_ during the *n*-th interpulse period, we break it into two portions: 1) pulse with nonzero laser intensity where the evolution of the FRET pair is described by the propagator 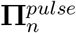 introduced shortly; 2) dark period with zero illumination intensity where the evolution of the FRET pair is described by the propagator 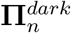 introduced shortly. Furthermore, depending on whether a photon is detected or not over the *n*-th interpulse period the propagators 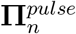 and 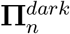 assume different forms.

First, when no photons are detected, we have

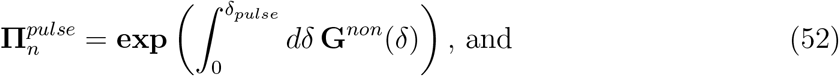

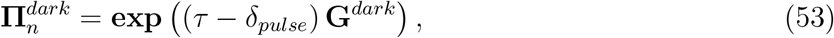

where the integration over the pulse period now involves a time dependent **G**^*non*^ due to temporal variations in λ_*ex*_(*t*). The integral in Eq. 52 is sometimes termed the excitation IRF though we will not use this convention here. For this reason, when we say IRF, we imply detection IRF alone. Additionally, **G**^*dark*^ is the same as **G**^*non*^ except for the excitation rate that is now set to zero due to lack of illumination. Finally, the propagator for an interpulse period with no photon detection can now be written as

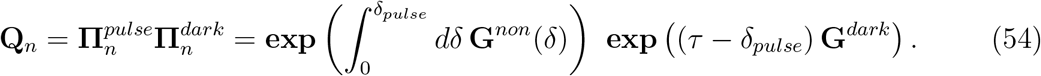

On the other hand, if a photon is detected sometime after a pulse (as in Fig. 3(a)), the pulse propagator remains as in Eq. 52. However, the propagator 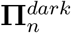 must now be modified to include a radiative generator matrix 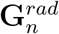 similar to Eq. 20

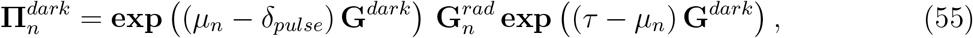

where *μ_n_* is the photon arrival time measured with respect to the *n*-th pulse (also termed microtime) as shown in Fig. 3(a). Here, the two exponential terms describe the evolution of the superstate before and after the photon detection during the dark period.

Moreover, we can construct the propagator for situations where a photon is detected during a pulse itself in a similar fashion. Here, the propagator 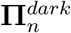 remains the same as in Eq. 53 but 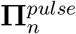 must now be modified to include the radiative generator matrix 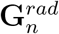 as

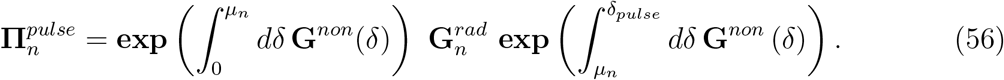

The propagators derived so far in this section assumed ideal detectors. We now describe a procedure to incorporate the IRF into this formulation. This is specially significant in accurate estimation of fluorophores’ lifetimes, which is commonly done in pulse illumination smFRET experiments. To incorporate the IRF, we follow the same procedure as in Sec. 2.4.3 and introduce convolution between the IRF function *f*(*ϵ*) and propagators above involving photon detections. That is, when there is a photon detected during the dark period, we modify the propagator 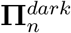 as

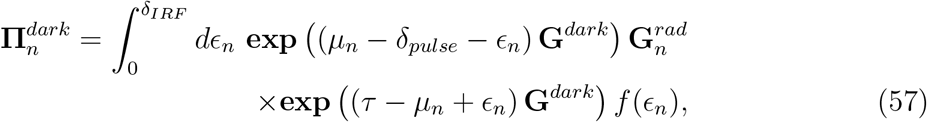

while the 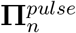 stays the same as in Eq. 52. Here, *ϵ_n_* is the stochastic delay in photon detection resulting from the IRF as shown in Fig. 3(b).

Moreover, when there is a photon detected during a pulse, the propagator 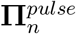 of Eq. 56 can be modified in a similar fashion to accommodate the IRF, while the propagator 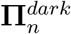 remains the same as in Eq. 53.

The propagators **Q**_*n*_ presented in this section involve integrals over large generator matrices that are analytically intractable and computationally expensive when considering large pulse numbers. Therefore, we follow a strategy similar to Sec. 2.3.8 for binned likelihood to approximate these propagators.

To reduce the complexity of the calculations, we start by making realistic approximations. Given the timescale separation between the interpulse period (typically tens of nanoseconds) and the system kinetics (typically of seconds timescale) in a pulsed illumination experiment, it is possible to approximate the system state trajectory as being constant during an interpulse period. In essence, rather than treating the system state trajectory as a continuous time process, we discretize the trajectory such that system transitions only occur at the beginning of each interpulse period. This allows us to separate the photophysical part of the generator matrix **G**_*ψ*_ in Eq. 4 from the portion describing the evolution of the system under study **G**_*σ*_ given in Eq. 3. Here, by contrast to the likelihood shown in Sec. 2.5.1 for pulsed illumination, we can now independently compute photophysical and system likelihood portions, as described below.

To derive the likelihood, we begin by writing the system state propagator during an interpulse period as

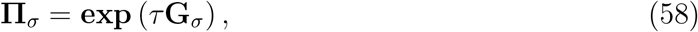

Furthermore, we must incorporate observations into these propagators by multiplying each system transition probability in **Π**_*σ*_, *π*_*σ_i_*→*σ_j_*_, with the observation probability if that transition had occurred. We organize these observation probabilities using our newly defined detection matrices 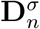 similar to Sec. 2.3.6, and write the modified propagators as

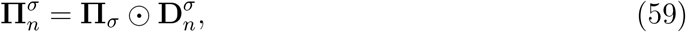

where ⊙ again represents the element-by-element product. Here, the elements of 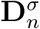 depend on the photophysical portion of the generator matrix **G**_*ψ*_ and their detailed derivations are shown in the third companion manuscript [64]. We note here that propagator matrix dimensions are now *M_σ_* × *M_σ_* making them computationally less expensive than in the continuous illumination case. Finally, the likelihood for the pulsed illuminated smFRET data with these new propagators reads

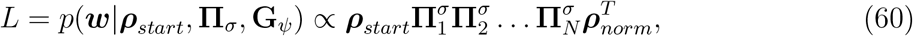

which, similar to the case of binned likelihood under continuous illumination (see Sec. 2.3.8), sums over all possible system state trajectories.

We will later use this likelihood to put forward an inverse model to learn transition probabilities (elements of **Π**_*σ*_) and photophysical transition rates appearing in **G**_*ψ*_.

### 2.6 Background Emissions

Here, we consider background photons registered by detectors from sources other than the labeled system under study [2]. The majority of background photons comprise ambient photons, photons from the illumination laser entering the detectors, and dark current (false photons registered by detectors) [22].

Due to the uniform laser intensity in the continuous illumination case, considered in this section, we may model all background photons using a single distribution from which waiting times are drawn. Often, such distributions are assumed (or verified) to be Exponential with fixed rates for each detection channel [65, 66]. Here we model the waiting time distribution for background photons arising from both origins as a single Exponential as is often the most common case. However, in the pulsed illumination case, laser source and the two other sources of background require different treatments due to nonuniform laser intensity. That is, the ambient photons and dark current are still modeled by an Exponential distribution though it is often further approximated as a Uniform distribution given that interpulse period if much shorter than the average background waiting time. The full formulation describing all background sources under pulsed illumination is provided in the third companion manuscript [64].

We now proceed to incorporate background into the likelihood under continuous illumination. We do so, as mentioned earlier, by assuming an Exponential distribution for the background, which effectively introduces new photophysical transitions into the model. As such, these transitions may be incorporated by expanding the full generator matrix **G** (described in Sec. 2.3) appearing in the likelihood, thereby leaving the structure of the likelihood itself intact, *c.f.*, Eq. 21.

To be clear, in constructing the new generator matrix, we treat background in each detection channel as if originating from fictitious independent emitters with constant emission rates (exponential waiting time). Furthermore, we assume that an emitter corresponding to channel *i* is a two state system with photophysical states denoted by

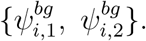

Here, each transition to the other state coincides with a photon emission with rate 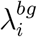. As such, the corresponding background generator matrix for channel *i* can now be written as

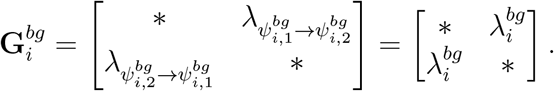

Since the background emitters for each channel are independent of each other, the expanded generator matrix **G** for the combined setup (system-FRET composite plus background) can now be computed. This can be achieved by combining the system-FRET composite state space and the background state spaces for all of the total *C* detection channels using Kronecker sums [67] as

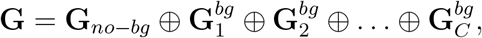

where the symbol ⊕ denotes the matrix Kronecker sum, and **G**_*no-b*_ represents previously shown generator matrices without any background transition rates.

The propagators needed to compute the likelihood can now be obtained by exponentiating the expanded generator matrix above as

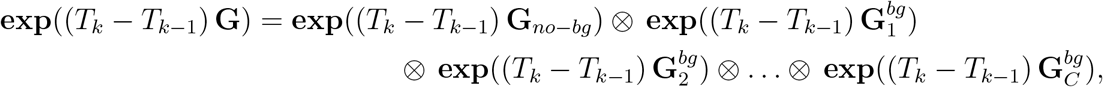

where the symbol ⊗ denotes the matrix Kronecker product (tensor product) [67].

Furthermore, the same detection matrices defined earlier to include only nonradiative transitions or only radiative transitions, and their generalization with crosstalk and detection efficiency, can be used to obtain nonradiative and radiative propagators, as shown in Sec. 2.3.6.

Consequently, as mentioned earlier, by contrast to incorporating the effects of dead time or IRF, addition of background sources do not entail any changes in the basic structure (arrangement of propagators) of the likelihood appearing in Eq. 21.

##### Example VI: Background

To provide a concrete example for background, we again return to our FRET pair with two system states. The background free full generator matrix for this system-FRET composite was provided in the example box in Sec. 2.3 as (in units of ms^−1^)

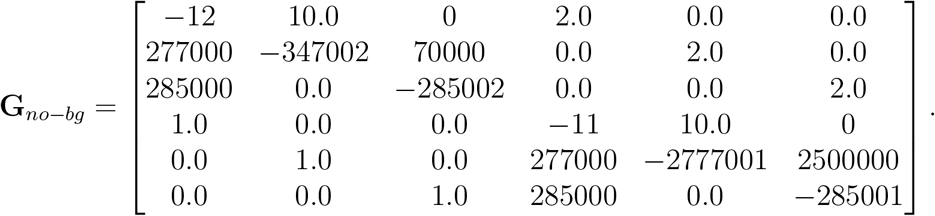

Here, we expand the above generator matrix to incorporate background photons entering two channels (i = 1, 2) at rates of 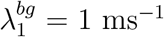 and 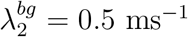. We do so by performing a Kronecker sum of **G**_*no-bg*_ with the following generator matrix for the background

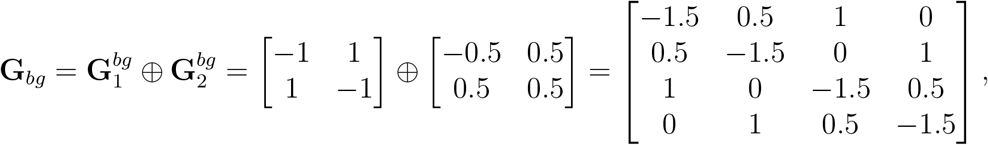

resulting in

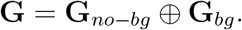

Here, **G** is a 24× 24 matrix and we do not include its explicit from.

### 2.7 Fluorophore Characteristics: Quantum Yield, Blinking, Photobleaching, and Direct Acceptor Excitation

As demonstrated for background in the previous section, to incorporate new photophysical transitions such as fluorophore blinking and photobleaching into the likelihood, we must modify the full generator matrix **G**. This can again be accomplished by adding extra photophysical states, relaxing nonradiatively, to the fluorophore model. These photophysical states can have long or short lifetimes depending on the specific photophysical phenomenon at hand. For example, donor photobleaching can be included by introducing a third donor photophysical state into the matrix of Eq. 4 without any escape transitions as follows

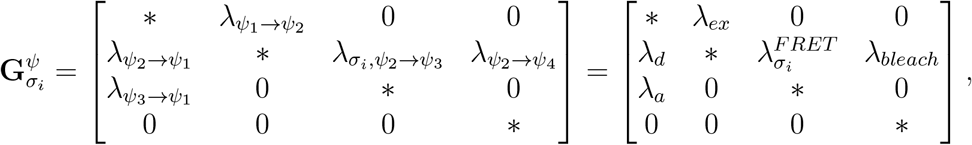

where *ψ*_1_ is the lowest energy combined photophysical state for the FRET labels, *ψ*_2_ represents the excited donor, *ψ*_3_ represents the excited acceptor, and *ψ*_4_ represents a photobleached donor, respectively. Additionally, λ_*d*_ and λ_*a*_ denote donor and acceptor relaxation rates, respectively, λ_*bleach*_ represents permanent loss of emission from the donor (photobleaching), and 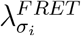 represents FRET transitions when the system is in its *i*-th system state.

Fluorophore blinking can be implemented similarly, except with a nonzero escape rate out of the new photophysical state, allowing the fluorophore to resume emission after some time [52, 68]. Here, assuming that the fluorophore cannot transition into the blinking photophysical state from the donor ground state results in the following generator matrix

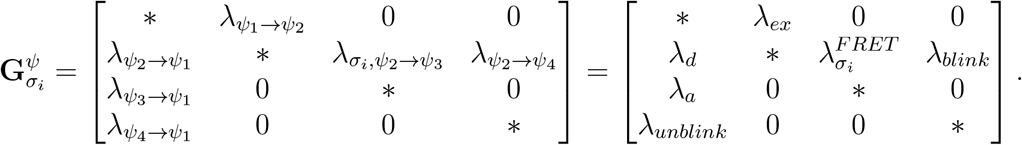

So far, we have ignored direct excitation of acceptor dyes in the likelihood model. This effect can also be incorporated into the likelihood by assigning a nonzero value to the transition rate λ*ψ*_1_→*ψ*_3_, that is,

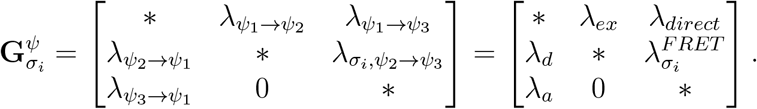

Other photophysical phenomena can also be incorporated into our likelihood by following the same procedure as above. Finally, just as when adding background, the structure of the likelihood (arrangement of the propagators) when treating photophysics (including adding the effect of direct acceptor excitation) stays the same as in Eq. 21.

### 2.8 Synthetic Data Generation

In the previous subsections, we described how to compute the likelihood, which is the sum of probabilities over all possible superstate trajectories that could give rise to the observations made by a detector, as demonstrated in Sec. 2. Here, we demonstrate how one such superstate trajectory can be simulated to produce synthetic photon arrival data using the Gillespie algorithm [69], as described in the next section followed by the addition of detector artefacts. We then use the generated data to test our BNP-FRET sampler.

#### 2.8.1 Gillespie and detector artefacts

The Gillespie algorithm generates two sets of random variables. The times at which superstates change (indexed 1 through *N*). These times occur anywhere along a continuous time grid. The next set of random variables are the states associated to the superstate preceding the time at which the superstate changes.

We designate the sequence of superstates

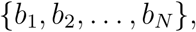

where *b_n_* ∈ {*ϕ*_1_, *ϕ*_2_,…, *ϕ_M_ϕ__*}. Here, unlike earlier in Sec. 2.3, the time index *n* on superstates *b_n_* is not on a regular temporal grid.

Now, to generate the superstate sequence above, we first randomly draw the first superstate, *b*_1_, from the set of possible superstates given their corresponding probabilities. Next, we draw the second superstate *b*_2_ of the sequence using the set of transition rates out of the first state with self-transitions excluded by construction. Now, after choosing *b*_2_, we generate the holding time *h*_1_ (the time spent in *b*_1_) from the Exponential distribution with rate constant associated with transitions *b*_1_ → *b*_2_. Finally, we repeat the two previous steps to sequentially generate the full sequence of superstates along with the corresponding holding times.

More formally, we generate a trajectory, by first sampling the initial superstate as

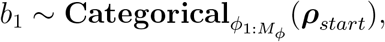

where ***ρ***_*start*_ is the initial probability vector and the Categorical distribution is the generalization of the Bernoulli distribution for more than two possible outcomes. The remaining superstates can now be sampled as

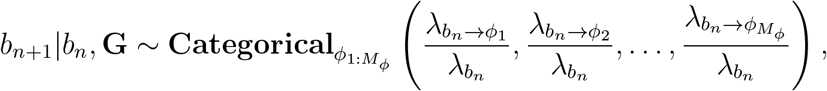

where λ_*b_n_*_ = ∑_*i*_ λ_*b_n_*→*ϕ_i_*_ is the escape rate for the superstate *b_n_* and rates for self-transitions are zero. The above equation reads as follows: “the superstate *b*_*n*+1_ is drawn (sampled) from a Categorical distribution given the superstate *b_n_* and the generator matrix **G**”.

Once the *n*-th superstate *b_n_* is chosen, the holding time *h_n_* (the time spent in *b*_n_) is sampled as follows

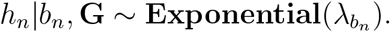

Finally, with ideal detectors, the detection channel *c_k_* is assigned deterministically to the *k*-th photon emitted at time 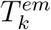, which can be computed by summing all the holding times preceding the corresponding radiative transition.

Furthermore, in the presence of detection effects, such as crosstalk, detection efficiency, and IRF, we must add to the stochastic output of the Gillespie simulation another layer of stochasticity originating from the measurement model. That is, we stochastically assign detection channel and detection times to an emitted photon, as described below.

In the presence of crosstalk and inefficient detectors, we choose the detection channel for the *k*-th photon emitted upon a radiative transition as

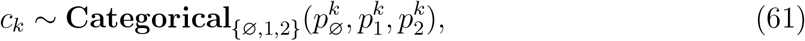

where 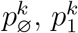 and 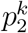, respectively, denote the probability of the photon going undetected, being detected in channel 1 and channel 2.

Moreover, in the presence of the IRF, we assign a stochastic delay *ϵ_k_*, sampled from a probability distribution *f*(*ϵ*), to the absolute photon emission time 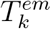. This results in the detection time, 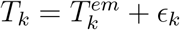, as registered by the timing hardware.

Additionally, when photophysical effects (such as blinking and photobleaching) and background are present, we can generate a superstate trajectory following the same procedure as above using the generator matrices **G** incorporating these effects as described in the previous sections.

Finally, we obtain our desired smFRET trace (see Fig. 4) consisting of photon arrival times *T*_1:*K*_ and detection channels *c*_1:*K*_ as

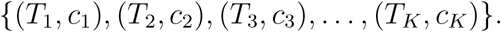

**Figure 4:**
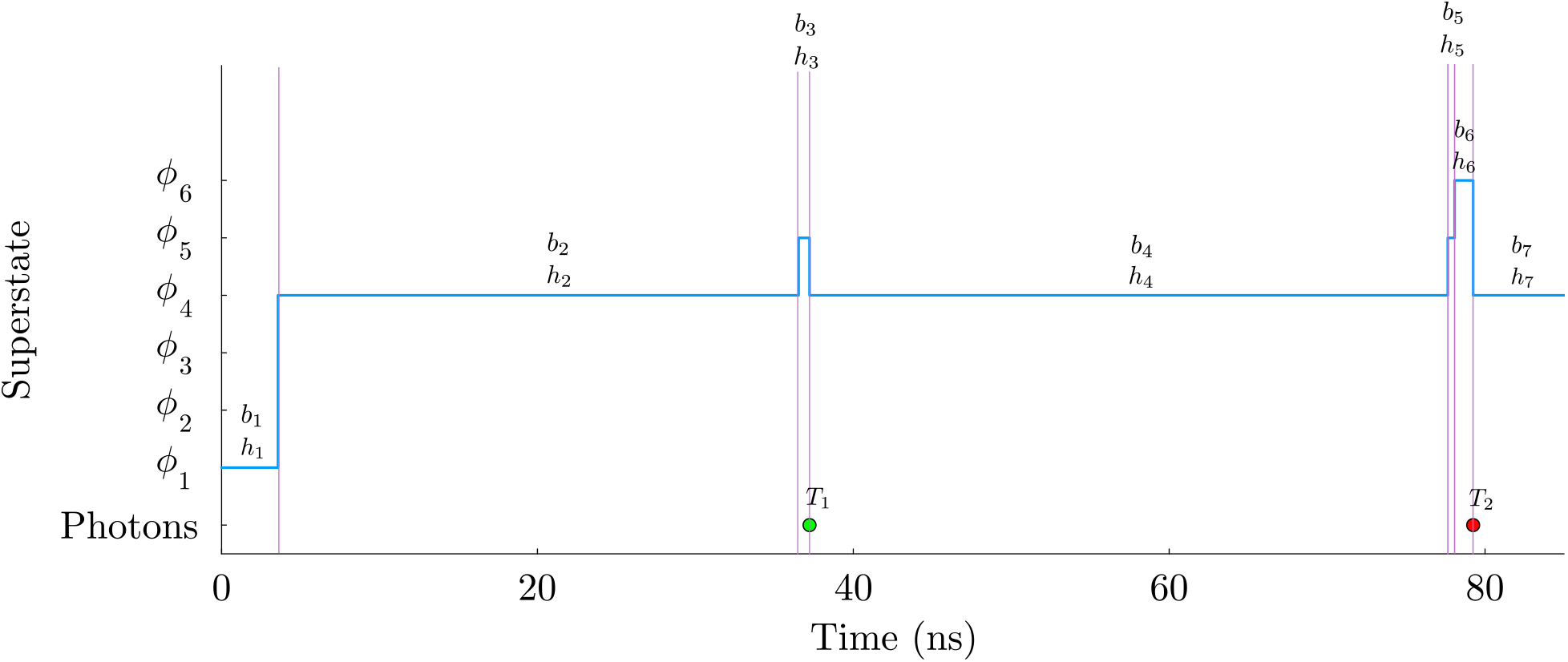
Simulated data. Here, we show a superstate trajectory (in blue) generated using Gillespie algorithm for a system-FRET composite with two system states *σ*_1:2_ and three photophysical states. Collectively, superstates ϕ_1:3_ correspond to photophysical states when the system resides in *σ*_1_ and superstates *ϕ*_4:6_ correspond to photophysical states when system resides in *σ*_2_. The pink vertical lines mark the time points where transitions between the superstates occur. The variables *b*_1:7_ and *h*_1:7_ between each set of vertical lines represent the superstates and associated holding times, respectively. The green and red dots show the photon detections at times *T*_1_ and *T*_2_ in the channels 1 and 2, respectively. The first photon is detected upon transition *b*_3_ → *b*_4_ (or *ϕ*_5_ → *ϕ*_4_), while the second photon is detected upon transition *b*_6_ → *b*_7_ (or *ϕ*_6_ → *ϕ*_4_). For this plot, we have used very fast system transition rates of λ_*σ*_1_→*σ*_2__ = 0.001ns^−1^ and λ_*σ*_1_→*σ*_2__ = 0.002ns^−1^ for demonstrative purposes only.

## 3 Inverse Strategy

Now, armed with the likelihood for different experimental setups and a means by which to generate synthetic data (or having experimental data at hand), we proceed to learn the parameters of interest. Assuming precalibrated detector parameters, these include transition rates that enter the generator matrix **G**, and elements of ***ρ***_*start*_. However, accurate estimation of the unknowns requires an inverse strategy capable of dealing with all existing sources of uncertainty in the problem such as photon’s stochasticity and detector noise. This naturally leads us to adopt a Bayesian inference framework where we employ Monte Carlo methods to learn distributions over the parameters.

We begin by defining the distribution of interest over the unknown parameters we wish to learn termed the posterior. The posterior is proportional to the product of the likelihood and prior distributions using Bayes’ rule as follows

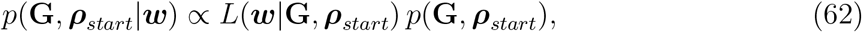

where the last term *p*(**G**, ***ρ***_*start*_) is the joint prior distribution over **G** and ***ρ***_*start*_ defined over the same domains as the parameters. The prior is often selected on the basis of computational convenience. The influence of the prior distribution on the posterior diminishes as more data is incorporated through the likelihood. Furthermore, the constant of proportionality is the inverse of the absolute probability of the collected data, 1/*p*(***w***), and can be safely ignored as generation of Monte Carlo samples only involves ratios of posterior distributions or likelihoods.

Additionally, the *ϵ^K^* factor in the likelihood first derived in Eq. 21 can be absorbed into the proportionality constant as it does not depend on any of the parameters of interest, resulting in the following expression for the posterior (in the absence of detector dead time and IRF for simplicity)

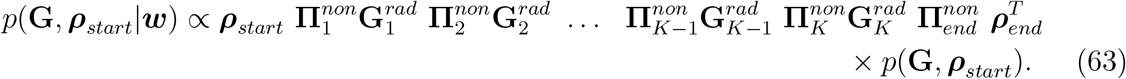

Next, assuming *a priori* that different transition rates are independent of each other and initial probabilities, we can simplify the prior as follows

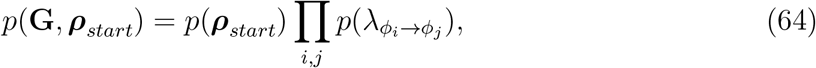

where we select the Dirichlet prior distribution over initial probabilities as this prior is conveniently defined over a domain where the probability vectors, drawn from it, sum to unity. That is,

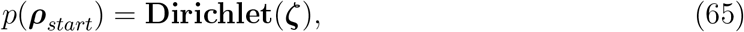

where the Dirichlet distribution is a multivariate generalization of the Beta distribution and ***ζ*** is a vector of the same size as the superstate space. Typically parameters of the prior are termed hyperparameters and as such ***ζ*** collects as many hyperparameters as its size.

Additionally, we select Gamma prior distributions for individual rates. That is,

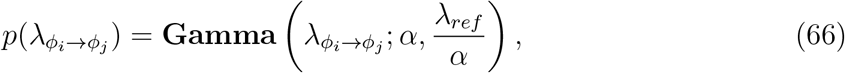

guaranteeing positive values. Here, *α* and λ_*ref*_ (a reference rate parameter) are hyperparameters of the Gamma prior. For simplicity, these hyperparameters are usually chosen (with appropriate units) such that the prior distributions are very broad, minimizing their influence on the posterior.

Furthermore, to reduce computational cost, the number of parameters we need to learn can be reduced by reasonably assuming the system was at steady-state immediately preceding the time at which the experiment began. That is, instead of sampling ***ρ***_*start*_ from the posterior, we compute ***ρ***_*start*_ by solving the time-independent master equation,

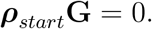

Therefore, the posterior in Eq. 63 now reduces to

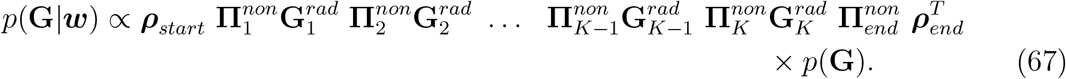

In the following subsections, we first describe a parametric inverse strategy, *i.e.*, assuming a known number of system states, for sampling parameters from the posterior distribution in Eq. 67 using Monte Carlo methods. Next, we generalize this inverse strategy to a nonparametric case where we also deduce the number of system states.

### 3.1 Parametric Sampler: BNP-FRET with Fixed Number of System States

Now with the posterior, Eq. 67, at hand and assuming steady-state ***ρ***_*start*_, here we illustrate a sampling scheme to deduce the transition rates of the generator matrix **G**.

As our posterior of Eq. 67 does not assume a standard form amenable to analytical calculations, we must iteratively draw numerical samples of the transition rates within **G** using Markov Chain Monte Carlo (MCMC) techniques. Specifically, we adopt a Gibbs algorithm to, sequentially and separately, generate samples for individual transition rates at each MCMC iteration. To do so, we first write the posterior of Eq. 67 using the chain rule as follows

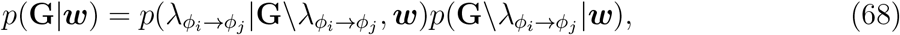

where the backslash after **G** indicates exclusion of the following rate parameters and ***w*** denotes the set of observations as introduced in Sec. 2.3.2. Here, the first term on the right hand side is the conditional posterior for the individual rate λ_*ϕ_i_*→*ϕ_j_*_. The second term is considered a constant in the corresponding Gibbs step as it does not depend on λ_*ϕ_i_*→*ϕ_j_*_. Moreover, following the same logic, the priors *p*(**G**\λ_*ϕ_i_*→*ϕ_j_*_.) (see Eq. 67) for the remaining rate parameters in the posterior on the left are also considered constant. Therefore, from Eqs. 67 & 68, we can write the conditional posterior for λ_*ϕ_i_*→*ϕ_j_*_ above as

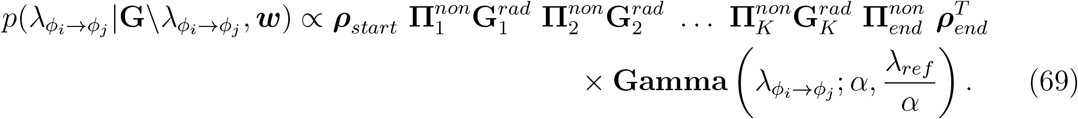

Just as with the posterior over all parameters, this conditional posterior shown above does not take a closed form allowing for direct sampling.

As such, we turn to the Metropolis-Hastings (MH) algorithm [70] to draw samples from this conditional posterior, where new samples are drawn from a proposal distribution *q* and accepted with probability

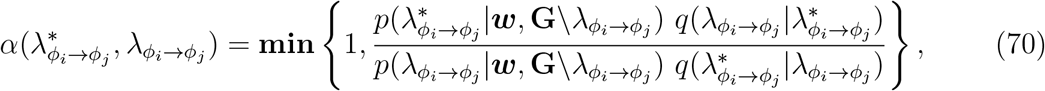

where the asterisk represents the proposed rate values from the proposal distribution *q*.

To construct an MCMC chain of samples, we begin by initializing the chain for each transition rate λ_*ϕ_i_*→*ϕ_j_*_, by random values drawn from the corresponding prior distributions. We then iteratively sweep the whole set of transition rates in each MCMC iteration by drawing new values from the proposal distribution *q*.

A computationally convenient choice for the proposal is a Normal distribution leading to a simpler acceptance probability in Eq. 70. This is due to its symmetry resulting in 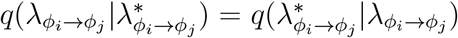. However, a Normal proposal distribution would allow negative transition rates naturally forbidden leading to rejection in the MH step and thus inefficient sampling. Therefore, it is convenient to propose new samples either drawn from a Gamma distribution or, as shown below, from a Normal distribution in logarithmic space to allow for exploration along the full real line as follows

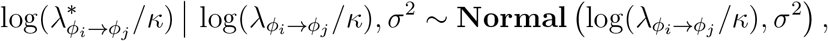

where *κ* =1 is an auxiliary parameter in the same units as λ_*ϕ_i_*→*ϕ_j_*_ introduced to obtain a dimensionless quantity within the logarithm.

The variable transformation above now requires introduction of Jacobian factors in the acceptance probability as follows

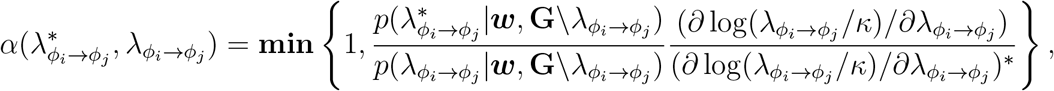

where the derivative terms represent the Jacobian and the proposal distributions are canceled by virtue of using a symmetric Normal distribution.

The acceptance probability above depends on the difference of the current and proposed values for a given transitions rate. In other words, smaller differences between the current and proposed values often lead to larger acceptance probabilities. This difference is determined by the covariance of the Normal proposal distribution σ^2^ which needs to be tuned for each rate individually to achieve optimal performance of the BNP-FRET sampler, or very approximately, one-fourth acceptance rate for the proposals [71].

This whole algorithm can now be summarized in the following pseudocode

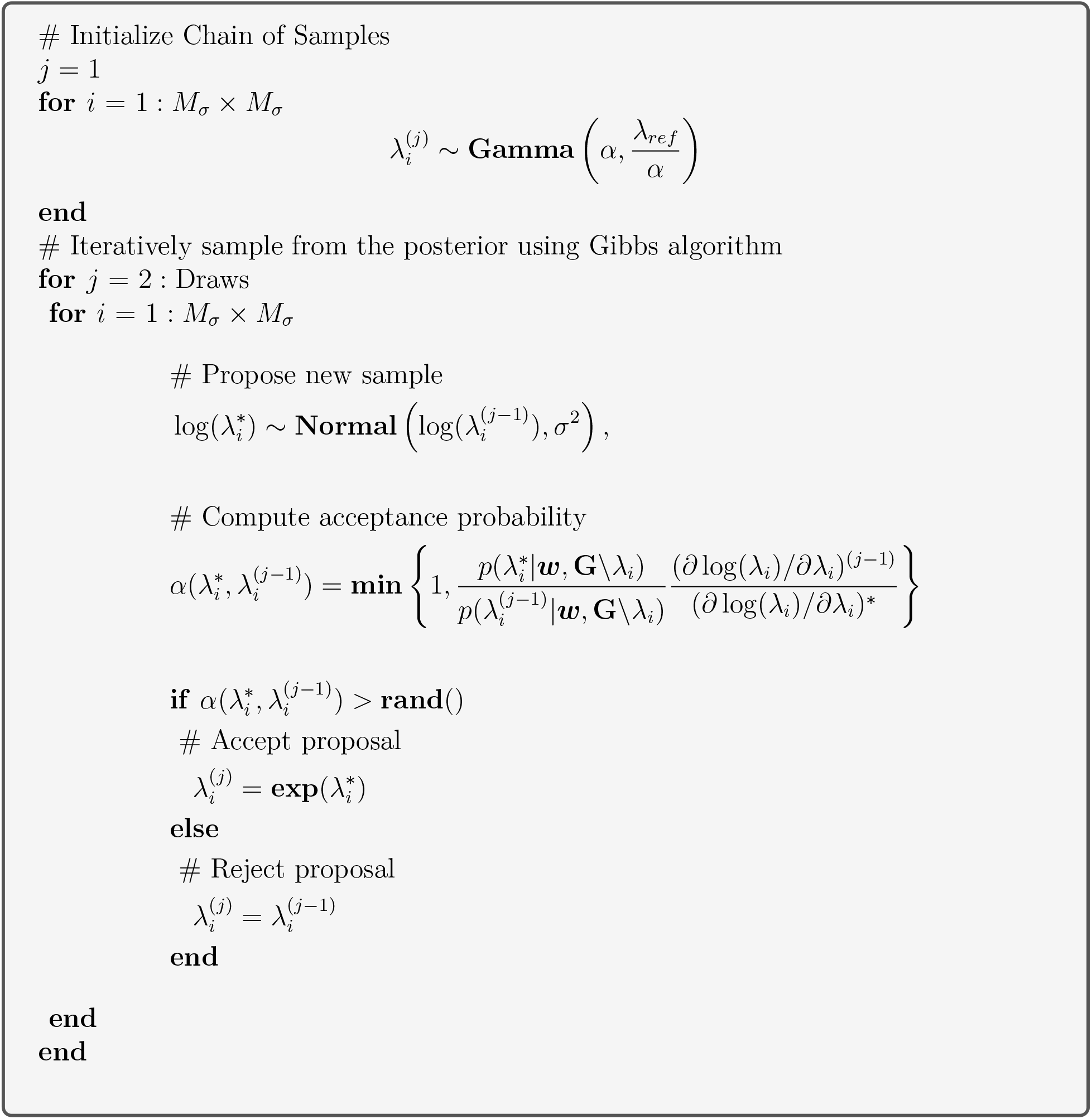

### 3.2 Nonparametrics: Predicting the Number of System States

After describing our inverse strategy for a known number of system states (*i.e.*, parametric inference), we turn to more realistic scenarios where we may not know the number of system states which, in turn, leads to an unknown number of superstates (*i.e.*, nonparametric inference). In the following subsections, we first describe the BNP framework for continuous illumination and then proceed to illustrate our BNP strategy under pulsed illumination. Such BNP frameworks introduced herein, eventually, provide us with distributions over the number of system states simultaneously, and self-consistently, with other model parameters.

#### 3.2.1 Bernoulli process for continuous illumination

The number of system states is often unknown and cannot *a priori* be set by hand. Therefore, in order to learn the states warranted by the data, we turn to the Bayesian nonparametric paradigm. That is, we first define an infinite-dimensional version of the generator matrix in Eq. 2 and multiply each of its elements by a Bernoulli random variable *b_i_* (also termed loads). These loads, indexed by *i*, allow us to turn on/off portions of the generator matrix associated with transitions between specific system states (including self-transitions). We can write the nonparametric generator matrix as follows

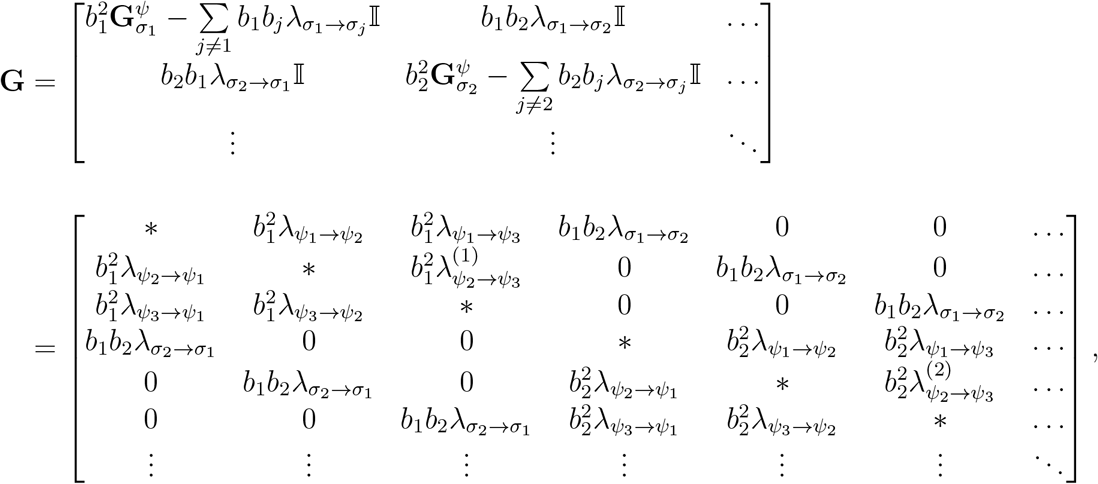

where a load value of 1 represents an “active” system state, while “inactive” system states (not warranted by the data) get a load value of 0. Here, there are two loads associated to every transition because there is a pair of states corresponding to each transition. Within this formalism, the number of active loads is the number of system states estimated by the BNP-FRET sampler. As before, * represents negative row-sums.

The full set of loads, **b** = {*b*_1_, *b*_2_,…,*b*_∞_}, now become quantities we wish to learn. In order to leverage Bayesian inference methods to learn the loads, the previously defined posterior distribution (Eq. 67) now reads as follows

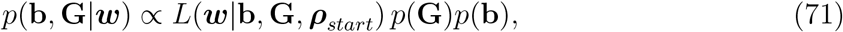

where the prior *p*(**b**) is Bernoulli while the remaining prior, *p*(**G**), can be assumed to be the same as in Eq. 66.

As in the case of the parametric BNP-FRET sampler presented in Sec. 3.1, we generate samples from this nonparametric posterior employing a similar Gibbs algorithm. To do so, we first initialize the MCMC chains of loads and rates by taking random values from their priors. Next, to construct the MCMC chains, we iteratively draw samples from the posterior in two steps: 1) sequentially sample all rates using the MH algorithm; then 2) loads by direct sampling, one-by-one from their corresponding conditional posteriors. Here, step (1) is identical to the parametric case of Sec. 3.1 and we only focus on the second step in what follows.

To sample the *i*-th load, the corresponding conditional posterior reads [72]

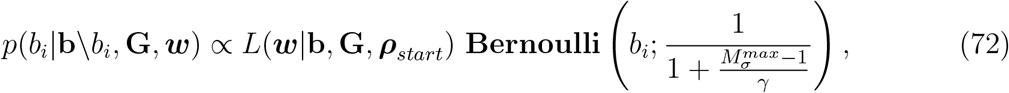

where the backslash after **b** indicates exclusion of the following load and 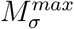 and *σ* are hyperparameters. Here 7 sets the *a priori* expected number of system states.

A note on the interpretation of 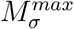 is in order. When dealing with nonparametrics, we nominally must consider an infinite set of loads and priors for these loads called Bernoulli process priors [73]. Samplers for such process priors are available though inefficient [74, 75]. However, for computational convenience, it is possible to introduce a large albeit finite number of loads set to 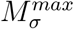 It can be shown that parameter inference are unaffected by this choice of cutoff [73, 76, 77] when setting the success probability to 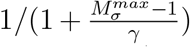 as in the Bernoulli distribution of Eq. 72. This is because such a choice forces the mean (expected number of system states) of the full prior on loads ∑_*i*_ *p*(*b_i_*) to be finite (= *σ*).

Since the conditional posterior in the equation above must be discrete and describes probabilities of the load being either active or inactive, it must itself follow a Bernoulli distribution with updated parameters

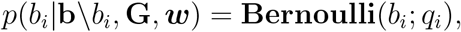

where

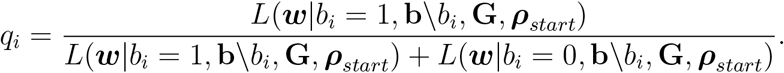

The Bernoulli form of this posterior allows direct sampling of the loads.

We will apply this method for synthetic and experimental data in the second companion manuscript of this series [78].

#### 3.2.2 iHMM methods for pulsed illumination

Under pulsed illumination, the Bernoulli process prior described earlier for continuous illumination can in principle be used as is to estimate the number of system states and the transition rates. However, in this section, we will describe a computationally cheaper inference strategy applicable to the simplified likelihood of Eq. 60 assuming system state transitions occurring only at the beginning of each pulse. The reduction in computational expense is achieved by directly learning the elements of the propagator **Π**_*σ*_ of Eq. 58, identical for all interpulse periods. In doing so, we learn transition probabilities for the system states instead of learning rates though we will continue learning rates for photophysical states. This avoids expensive matrix exponentials for potentially large system state numbers required for computing the propagators under continuous illumination.

Now, to infer the transition probabilities in **∑**_*σ*_, the dimensions of which are unknown owing to an unknown number of system states, as well as transition rates among the photophysical states (elements of **G**_*ψ*_ in Eq. 4), and initial probabilities, we must place suitable priors on these parameters yielding the following posterior

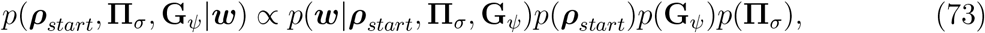

where we have immediately written the joint prior as a product prior over ***ρ***_start_, **G**_*ψ*_, and **Π**_*σ*_. Next, for ***ρ***_*start*_ and **G**_*ψ*_ we use the same priors as in Eqs. 65-66. However, as the number of system states is unknown, **Π**_*σ*_ requires special treatment. To learn **Π**_*σ*_, it is convenient to adopt the infinite HMM (iHMM) [41, 48] due to the discrete nature of system state transitions over time.

As the name suggests, the iHMM leverages infinite system state spaces (*M_σ_* → ∞ in Eq. 3) similar to the Bernoulli process prior of Sec. 3.2.1. However, unlike the Bernoulli process, all system states remain permanently active. The primary goal of an iHMM is then to infer transition probabilities between system states, some of which, not warranted by the data, remain very small and set by the (nonparametric) prior that we turn to shortly. Thus the effective number of system states can be enumerated from the most frequently visited system states over the course of a learned trajectory.

Within this iHMM framework, we place an infinite dimensional version of the Dirichlet prior, termed the Dirichlet process prior [41, 48, 79], as priors over each row of the propagator **Π**_*σ*_. That is,

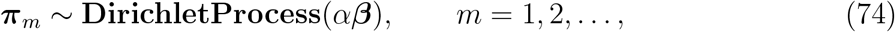

where ***π***_*m*_ is the mth row of **Π**_*σ*_. Here, the hyperparameters of the Dirichlet process prior include the concentration parameter *α* which determines the sparsity of the ***π***_*m*_ and the hyper parameter ***β*** which is a probability vector, also known as base distribution. Together *αβ* are related to the *ζ* introduced earlier for the (finite) Dirichlet distribution of Eq. 65.

Next, as the base distribution itself is unknown and all transitions out of each state should be likely to revisit the same set of states, we must place the same base distribution on all Dirichlet process priors placed on the rows of the propagator. To sample this unique base, we again choose a Dirichlet process prior [41, 80–82], that is,

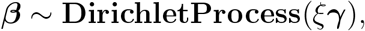

where we may set *ξ* = 1 and *σ* is a vector of hyperparameters of size *M_σ_*.

Now, to deduce the unknown parameters, we need to draw samples from the posterior in Eq. 73. However, due to the nonanalytical form of the posterior we cannot jointly sample our posterior. Thus, as before, we adopt a Gibbs sampling strategy to sequentially and separately draw samples for each parameter. Here, we only illustrate our Gibbs sampling step for the transition probabilities ***π***_*m*_. Our Gibbs steps for the remaining parameters are similar to Sec. 3.1. The complete procedure is described in the third companion manuscript [64].

Similar to the Bernoulli process prior, there are two common approaches to draw samples within the iHMM framework: slice sampling using the exact Dirichlet process prior and finite truncation [41, 48, 83, 84]. Just as before for the case of continuous illumination, we truncate the Dirichlet process prior to a finite Dirichlet distribution and fix its dimensionality to a finite (albeit large) number which we set to 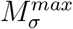 to improve the sampling. It can then be shown that for large enough 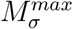 the number of system states visited becomes independent of 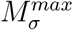 [72].

As before, to numerically sample the transition probabilities ***π***_*m*_ from our full posterior in Eq. 73 through MCMC, we choose our initial samples from the priors

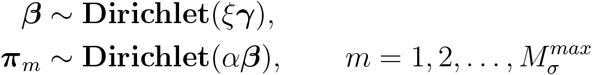

where we chose elements of *γ* to be 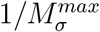 to ascribe similar weights across the state space *a priori*.

### 3.3 Likelihood Computation in Practice

As shown in Sec. 2.5.1, the likelihood typically takes the following generic form

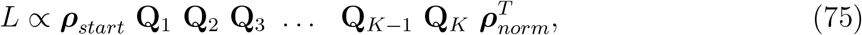

where **Q**_*i*_ are matrices whose exact form depends on which effects we incorporate into our likelihood. Computing this last expression would typically lead to underflow as likelihood values quickly drop below floating-point precision.

For this reason, it is convenient to introduce the logarithm of this likelihood. In order to derive the logarithm of the likelihood of Eq. 75, we rewrite the likelihood as a product of multiple terms as follows

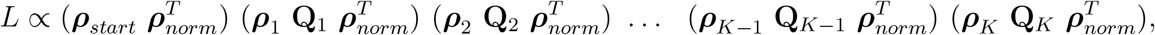

where ***ρ***_*i*_ are the normalized probability vectors given by the following recursive formula

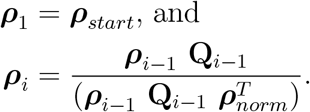

Now, using the recursion relation above, the log-likelihood can be written as

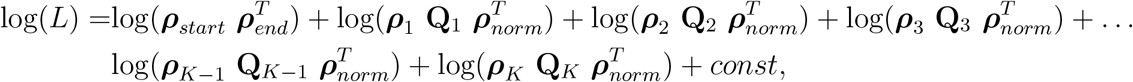

where const is a constant.

Note that 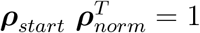. The pseudocode to compute the log-likelihood is as follows

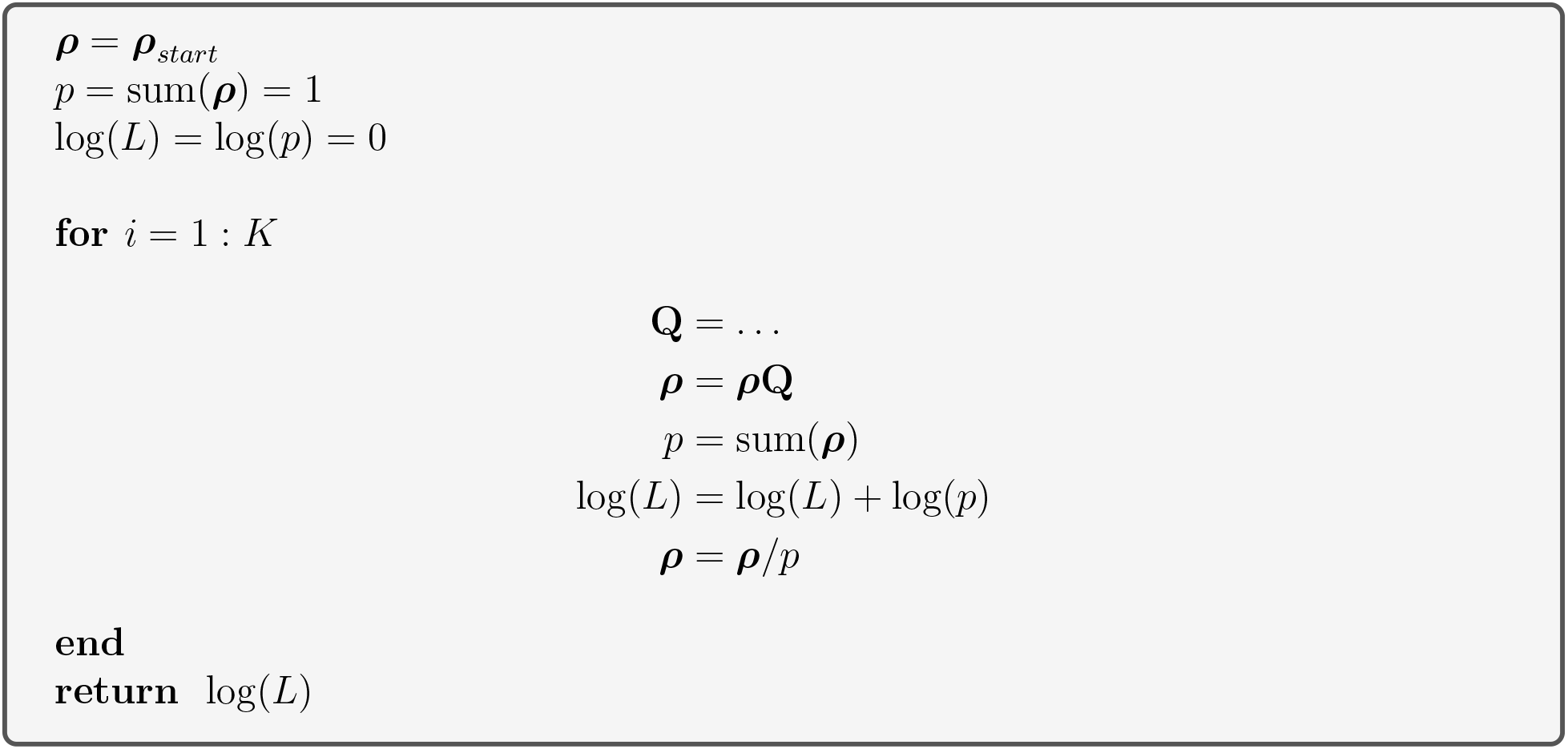

## 4 Results

In this section, we present results using our BNP-FRET sampler described above. Specifically, here we benchmark the parametric (*i.e.*, fixed number of system states) version of our sampler using synthetic data, while the two subsequent manuscripts [64, 78] focus on the nonparametric (*i.e.*, unknown number of system states) analysis of experimental data.

For simplicity alone, we begin by analyzing data from an idealized system with two system states using different photon budgets and excitation rates. Next, we consider more realistic examples incorporating the following one at a time: 1) crosstalk and detection efficiency; 2) background emission; 3) IRF; and then 4) end with a brief discussion on the unknown number of system states. We demonstrate when these features become relevant, as well as the overall robustness and generality of the BNP-FRET sampler.

For now, we assume continuous illumination for all parametric examples and use the following priors for the analyses. The prior used for the FRET rates are

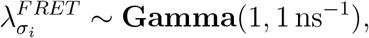

and use the following prior over the system transition rates

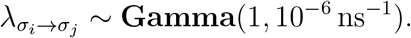

As discussed earlier in Sec. 3.1, it is more convenient to work within logarithmic space where we use the following proposal distributions to update the parameter values

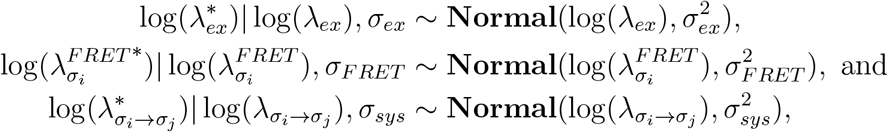

where * denotes proposed rates and where it is understood that all rates appearing in the logarithm have been divided through by a unit constant in order for the argument of the logarithm to remain dimensionless.

For efficient exploration of the parameter space by the BNP-FRET sampler and upon extensive experimentation with acceptance ratios, we found it prudent to alternate between two sets of variances, 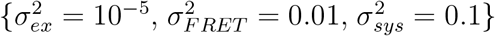 and 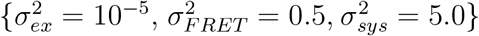 to generate an MCMC chain. This ensures that we propose samples of different orders of magnitude. As an intuitive guide, the more data we have, the sharper we expect our posterior over our rates to be and thus, the smaller both variances should be in our proposal distributions.

In the examples presented in the next few subsections, for computational simplicity, we fix the escape rates for the donor and acceptor excited photophysical states as well as the background rates for each detection channel in our simulations, as they can be precalibrated from experiments.

### 4.1 Parametric Examples

#### 4.1.1 Photon budget and excitation rate

Here, we perform Bayesian analysis on synthetically generated data (as described in Sec. 2.8) for the simplest case where the number of system states is an input to the BNP-FRET sampler. To generate data, we use the following generator matrix

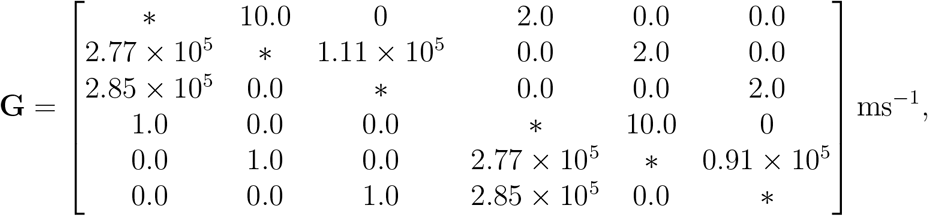

where the elements are motivated from real experiments [85]. Using this generator matrix, we generated a superstate trajectory as described in Sec. 2.8. We analyzed 430000 photons from the generated data using our BNP-FRET sampler. The resulting posterior distribution for transitions between system states and FRET efficiencies (computed as 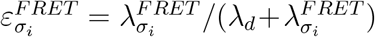 for the *i*-th system state) is shown in Fig. 5. As we will see for all examples, the finiteness of data always leads to some error as evident from the slight offset of the peaks of the distribution from the ground truth.

**Figure 5:**
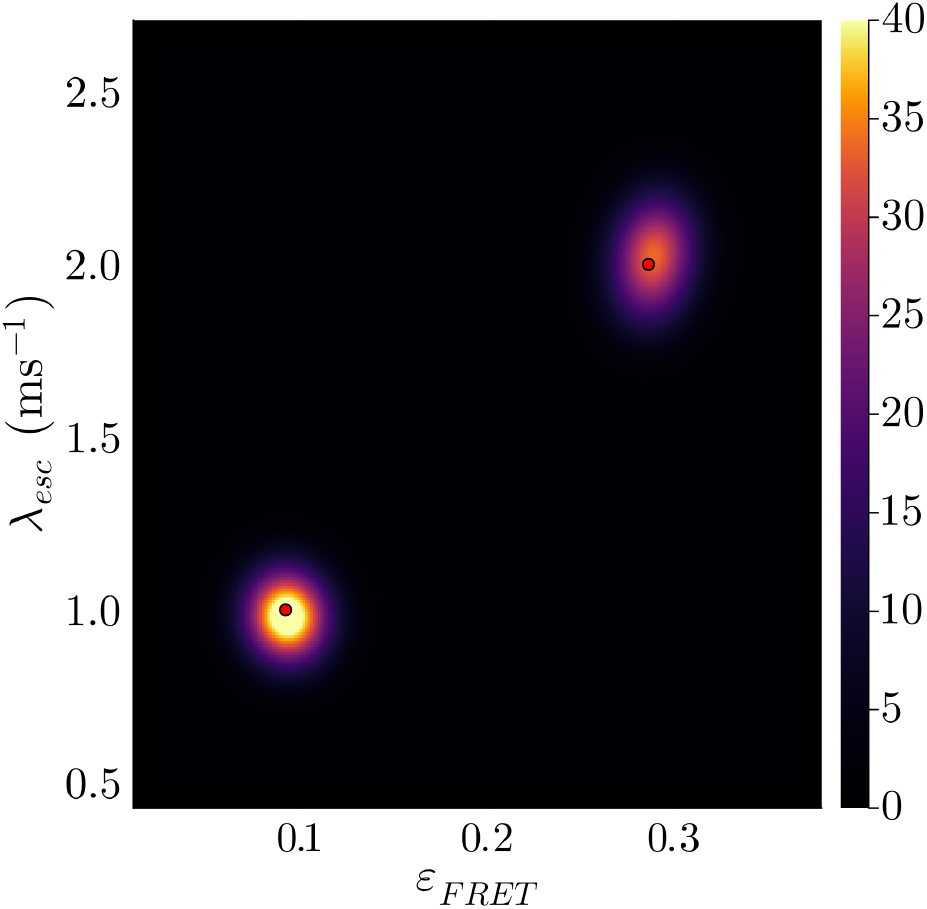
Learned bivariate posterior for the system state escape rates λ_*esc*_ and FRET efficiencies *ε_FRET_* given synthetic data. To produce this plot, we analyzed synthetic data generated using an excitation rate of λ_*ex*_ = 10 ms^-1^, and escape rates λ_*esc*_ = 1 & 2 ms^-1^ and FRET efficiencies of 0.09 & 0.29 for the two system states, respectively. The ground truth is shown with red dots. The FRET efficiencies estimated by our sampler are 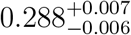 and 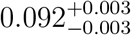. Furthermore, predicted escape rates are 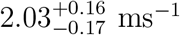 and 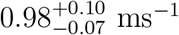. The small bias away from the ground truth is due to the finiteness of data. We have smoothed the distributions, for illustrative purposes only, using kernel density estimation (KDE) available through the Julia Plots package.

The effects of limited photon budget become significant especially when system kinetics occur across multiple timescales with the most photon starved state characterized by the largest escape rate. In this case, it is useful to quantify how many photons are typically required to assess any escape rate (with the fastest setting the lower photon count bound needed) to obtain below 15% error in parameter estimates.

To quantify the number of photons, ignoring background and detector effects, we define a dimensionless quantity which we call the “photon budget index” predicting the photon budget needed to accurately estimate the transition rates in the model as

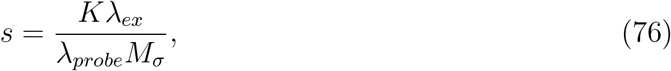

where *K* is the total number of photons in a single photon smFRET trace (photon budget), λ_*ex*_ is the excitation rate, λ_*probe*_ represents the escape rate (timescale) that we want to probe, and *M_σ_* is the number of system states. The parameters in the numerator control the amount of data available and the temporal resolution. On the other hand, the parameters in the denominator are the properties of the system under investigation and represent the required resolution.

From experimentation, we have found a photon budget index of approximately 10^6^ to be a safe lower threshold for keeping errors below 15% (this error cutoff is a user choice) in parameter estimates. In the simple parametric example above, we have *K* = 4.3 × 10^5^, λ_*ex*_ = 10 ms^-1^, and the fastest transition that we want to probe is λ_*probe*_ = 2 ms^-1^, and *M_σ_* = 2, which corresponds to a photon budget index of 1.08 × 10^7^. In Fig. 6, we also demonstrate the reduction in errors (confidence interval size) for parameters of the same system as the photon budget is increased from 12500 photons to 400000 photons. For each of those cases, we used 9000 MCMC samples to compute statistical metrics such as quantiles.

**Figure 6:**
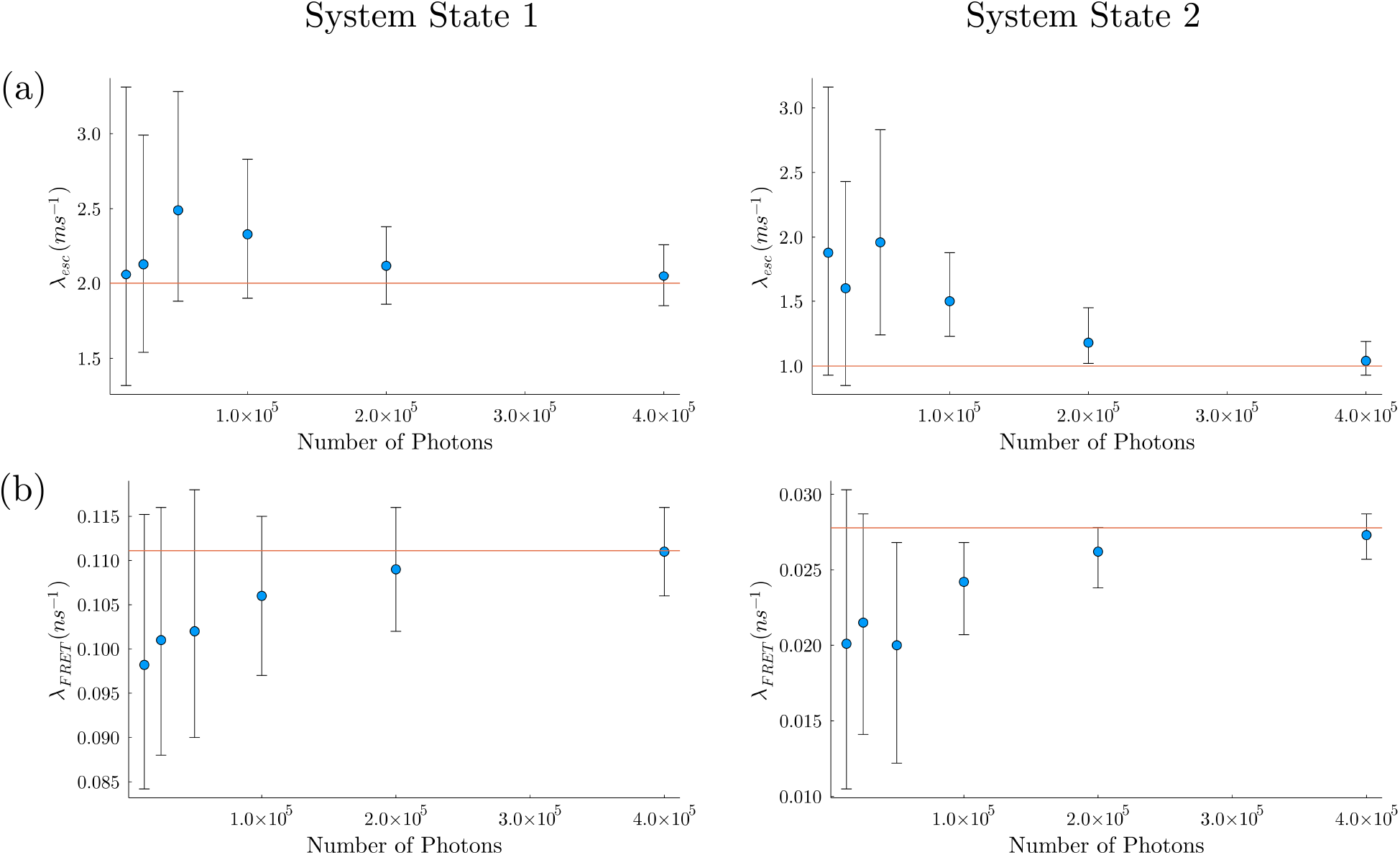
System and FRET transition rates as functions of the number of photons used for analysis. To produce these plots using the same kinetic parameters as in Fig. 5. Next, we analyzed the data considering only the first 12500 photons and then increased the photon budget by a factor of two for each subsequent analysis. Furthermore, we generated 9000 MCMC samples for each analysis to compute statistical quantities. In panel (a), we show two plots corresponding to the two system transition rates (λ_*esc*_). The blue dots represent the median values (50% quantile), and the ends of the attached confidence intervals represent 5% and 95% quantiles. The ground truths are shown with red horizontal lines. We show similar plots for FRET transition rates (λ_*FRET*_) in panel (b). In all of the plots, we see that as the photon budget is increased, the confidence intervals shrink (the posterior gets narrower/sharper). With a budget of 400000 photons, the confidence intervals represent less than 10% error in the estimates and contain the ground truths in all of the plots.

We further investigate the effect of another quantity which appears in the photon budget index, that is, excitation rate on the parameter estimates. To do so, we generate three new synthetic datasets, each containing ≈ 670000 photons, using the same excitation rate of 10 ms^−1^, and FRET efficiencies of 0.28 and 0.09 for the two system states, respectively, as before. However, the kinetics differ across these datasets so that they have system state transition rates well below, equal to, and well above the excitation rate. As such, for the first dataset, we probe slower kinetics compared to the excitation rate with system state transition rates set at 0.1 ms^−1^. In the next two datasets, the molecule changes system states at a much faster rates of 10 ms^−1^ and 1000 ms^−1^, respectively.

The results obtained for these FRET traces using our Bayesian methods are shown in Fig. 7. The bias in the posterior away from the ground truth increases as faster kinetics are probed in Fig. 7 from left to right. The results for the case with the fastest transition rates of 1000 ms^-1^ in Fig. 7(c) show a marked deterioration of the predictions, as the information content is not sufficient to separate the two FRET efficiencies resulting in estimated values close to the average of the ground truth values (≈ 0.185). This lack of information is also reflected in the uncertainties corresponding to each escape rate as shown in Fig. 7. Moreover, the predicted transition rates are of the same order as the excitation rate itself due to lack of temporal resolution available to probe such fast kinetics.

**Figure 7:**
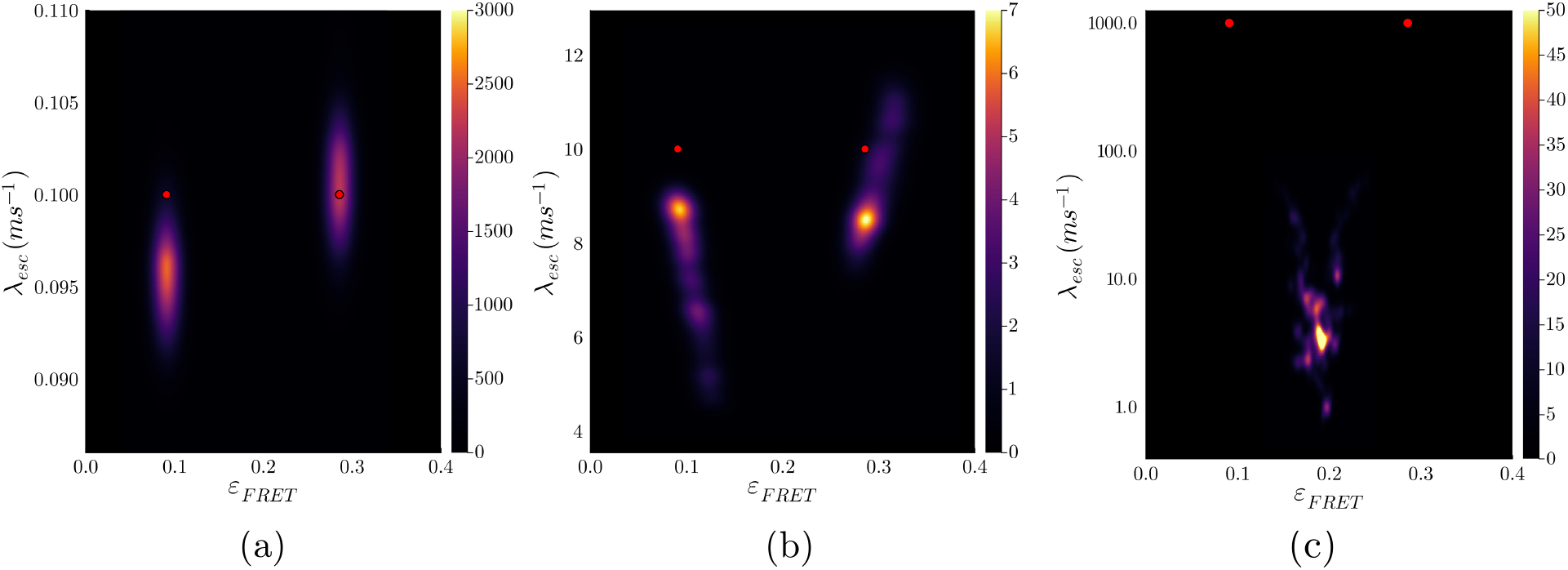
Learned bivariate posterior for the system state escape rates λ_*esc*_ (logscale in panel (c)) and FRET efficiencies *ε_FRET_* from synthetic data. For all synthetic smFRET traces, we use an excitation rate of 10 ms^-1^ and FRET efficiences of 0.29 and 0.09 for the two system states, respectively. The three panels correspond to different timescales being probed with transitions rates: (a) 0.1 ms^-1^; (b) 10 ms^-1^; and (c) 1000 ms^-1^. The ground truth values are shown with red dots. The bias in the parameter estimates increases as faster kinetics are probed, demonstrating deterioration of the information content of the collected data resulting in expectedly poor estimation assuming a fixed photon budget of 670000. This can also be seen quantitatively by calculating the confidence intervals reported below for each case. The FRET efficiencies estimated by our sampler for the slowest case in panel (a) are 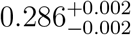 and 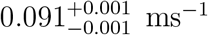, and the corresponding escape rates are 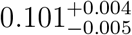 and 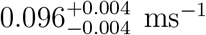. For the intermediate case in panel (b), FRET efficiencies estimated by our sampler are 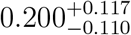 and 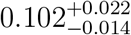, and predicted escape rates are 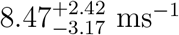 and 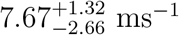. For the fastest case in panel (c), FRET efficiencies estimated by our sampler are 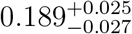 and 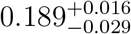, and predicted escape rates are 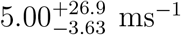 and 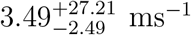. Poorer confidence intervals for larger escape rates reflect larger uncertainty due to lack of information.

To conclude, the excitation rate used to collect smFRET data and the total number of photons available determine the amount of information needed to resolve transitions among system states. As such, the ability of Bayesian methods to naturally propagate error from finiteness of information into parameter estimates make them indispensable tools for quantitative smFRET data analysis. This is by contrast to maximum likelihood-based methods which provide only inaccurate point estimates on account of limited data.

#### 4.1.2 An example with crosstalk

Here, we demonstrate how our method handles cases when significant crosstalk is present. To show this, we use the same dynamical parameters and photon budget as in the previous subsection for generating synthetic data but allow 5% of the donor photons to be stochastically detected by the acceptor channel. We then analyze the data with two versions of our method, one that incorporates crosstalk and one that ignores it altogether. Our results show that neglecting crosstalk necessarily leads to artefactually higher FRET efficiency estimates. This is clearly seen in Fig. 8(a). As expected, incorporation of crosstalk into the likelihood as shown in Sec. 2.4.1 results in a smaller bias. In this case, both ground truths falls within the range of posteriors for the corrected model; see Fig. 8(b). Furthermore, as shown in Fig. 9 top panels, we note that donor crosstalk again results in overestimation in FRET efficiencies. However, when we correct for crosstalk, our BNP-FRET sampler starts learning FRET efficiencies with ground truths falling within the range of 95% confidence intervals; Fig. 9 bottom panels. As expected, our simulations in Fig. 9 also show that uncertainty increases with increasing crosstalk and parameter estimation fails for crosstalk values beyond 60%.

**Figure 8:**
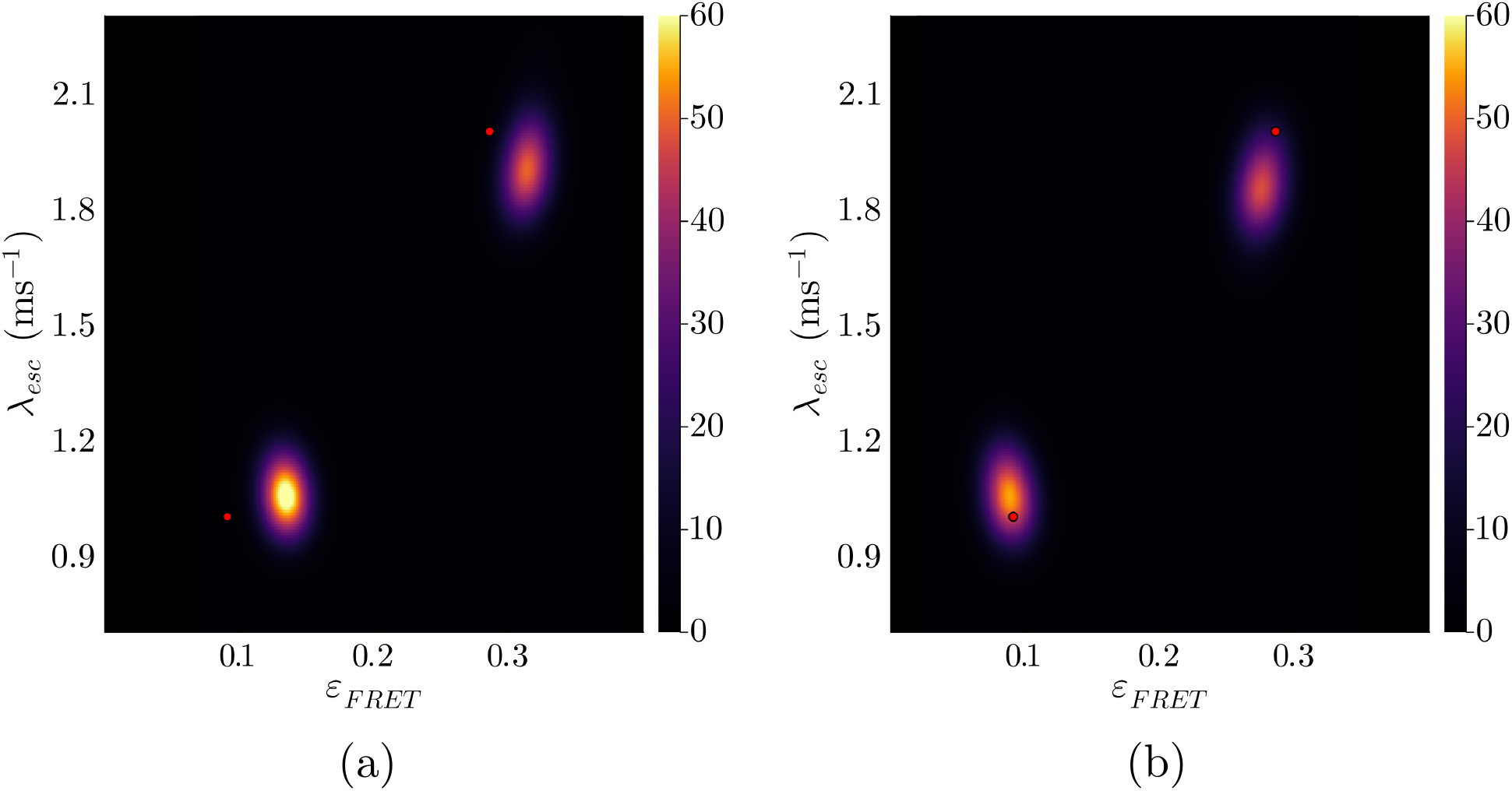
Learned bivariate posterior for the system state escape rates λ_*esc*_ and FRET efficiencies *ε_FRET_* for synthetic data with crosstalk. The ground truth is shown with red dots. In panel (a), we show the learned posterior using the model that doesn’t correct for cross talk consistently shows deviation from the ground truth with higher FRET efficiency estimates on account of more donor photons being detected as acceptor photons. The FRET efficiencies estimated by our sampler for this case are 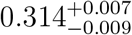 and 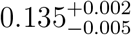, and the predicted escape rates are 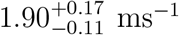 and 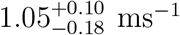. For the corrected case shown in panel (b), FRET efficiencies estimated by our sampler are 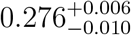 and 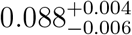. Furthermore, predicted escape rates are 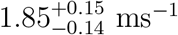 and 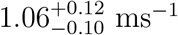. The corrected model mitigates this bias as demonstrated by the posterior in panel (b).

**Figure 9:**
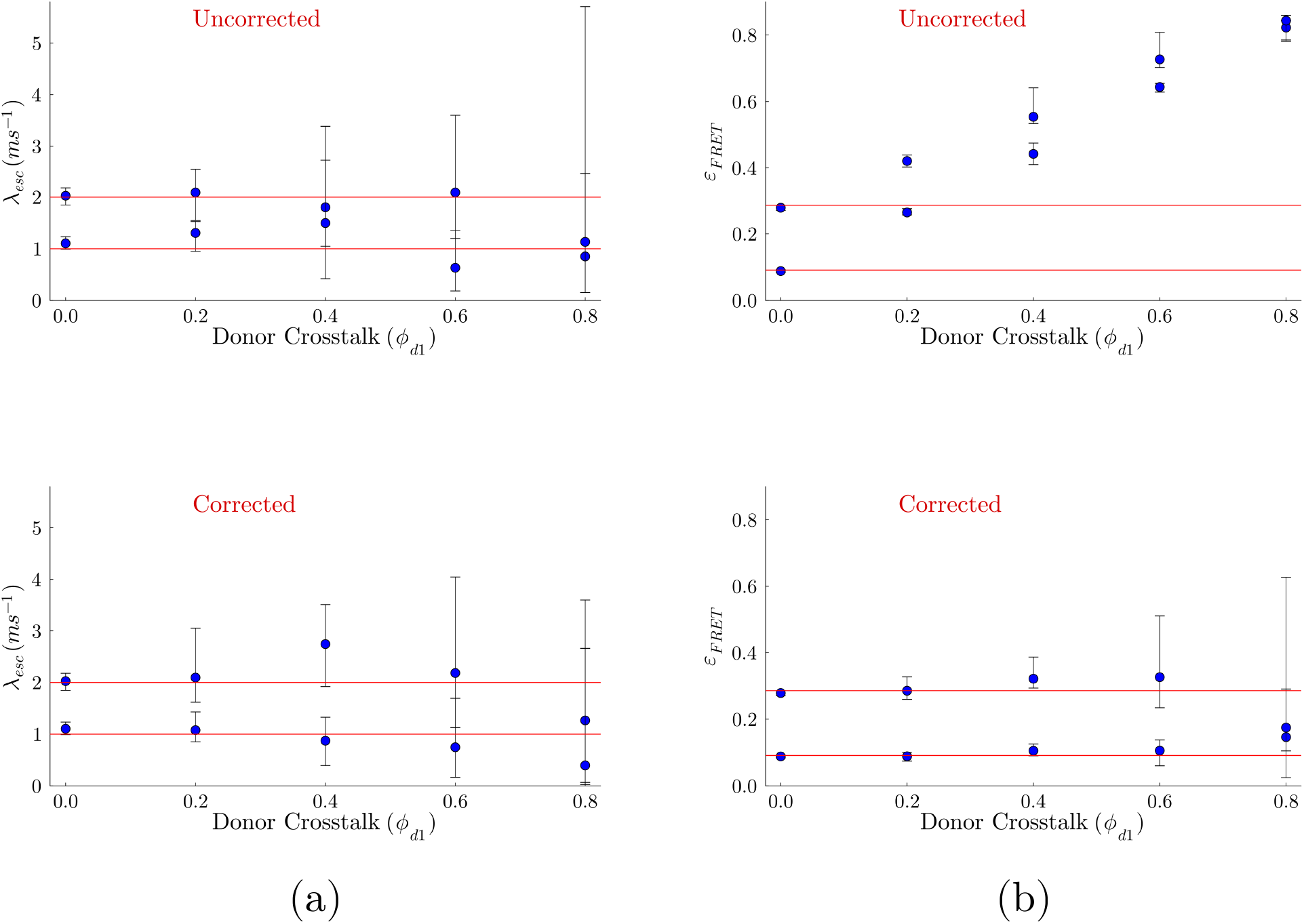
System transition rates λ_*esc*_ and FRET efficiencies *ε_FRET_* as functions of increasing donor crosstalk probability *ϕ*_*d*1_. To produce these plots, we generated synthetic data with excitation and escape rates as Fig. 5. In each plot, the blue dots represent the median values (50% quantile), and the ends of the attached confidence intervals represent the 5% and 95% quantiles. Furthermore, the ground truths are shown with red horizontal lines. In panel (a), our two plots show system transition rates estimated by the BNP-FRET sampler when corrected and uncorrected for crosstalk. We show similar plots for FRET efficiencies in panel (b). In all plots, we see that as donor crosstalk is increased, the confidence intervals grow (the posterior gets wider) and the estimates become unreliable after *ϕ*_*d*1_ > 0.6. Additionally, as expected, if uncorrected for, the FRET efficiencies start to merge with increasing crosstalk due to most photons being detected in acceptor channel (labeled 1).

#### 4.1.3 An example with background emissions

In section 2.6, we had shown a way to include background emissions in the forward model. For the current example, we again choose the same kinetic parameters for the system and the FRET pair as in Fig. 5 but now some of the photons come from background sources with rates 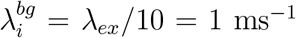 for the *i*-th channel. Addition of a uniform background would again lead to higher FRET efficiency estimates due to excess photons detected in each channel, if left uncorrected in the model, as can be seen in Fig 10(a) and Fig. 11 top panels. By comparison to the uncorrected method, our results migrate toward the ground truth when analyzed with the full method; see Fig 10(b) and Fig. 11 bottom panels. Furthermore, as shown in Fig. 11, when background photons account for more than approximately 40% of detected photons, relative uncertainties in estimated transition rates become larger than 25% indicating unreliable results.

**Figure 10:**
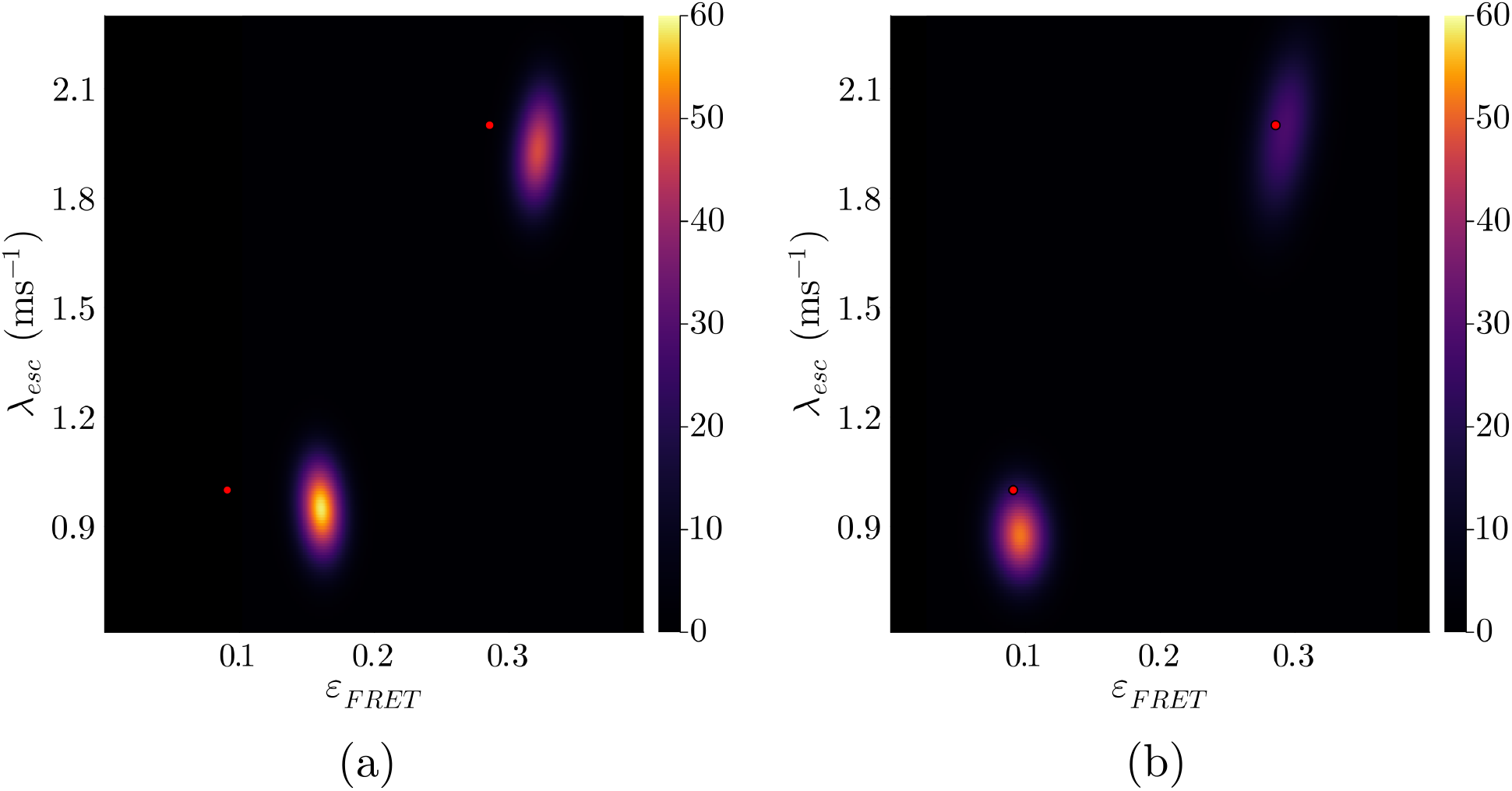
Learned bivariate posterior for the system state escape rates λ_*esc*_ and FRET efficiencies *ε_FRET_* given synthetic data with background emissions. The ground truth is shown with red dots. The learned posterior distribution using the model that doesn’t correct for background emissions (panel (a)) consistently shows bias away from the ground truth with higher estimates for FRET efficiencies on account of extra background photons. The FRET efficiencies estimated by our sampler for this uncorrected case are 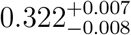 and 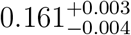, and the predicted escape rates are 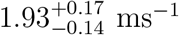 and 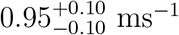. The corrected model mitigates this bias as demonstrated by the posterior in panel (b) as demonstrated by learned FRET efficiencies of 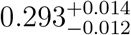 and 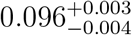. Furthermore, predicted escape rates are 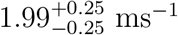 and 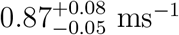.

**Figure 11:**
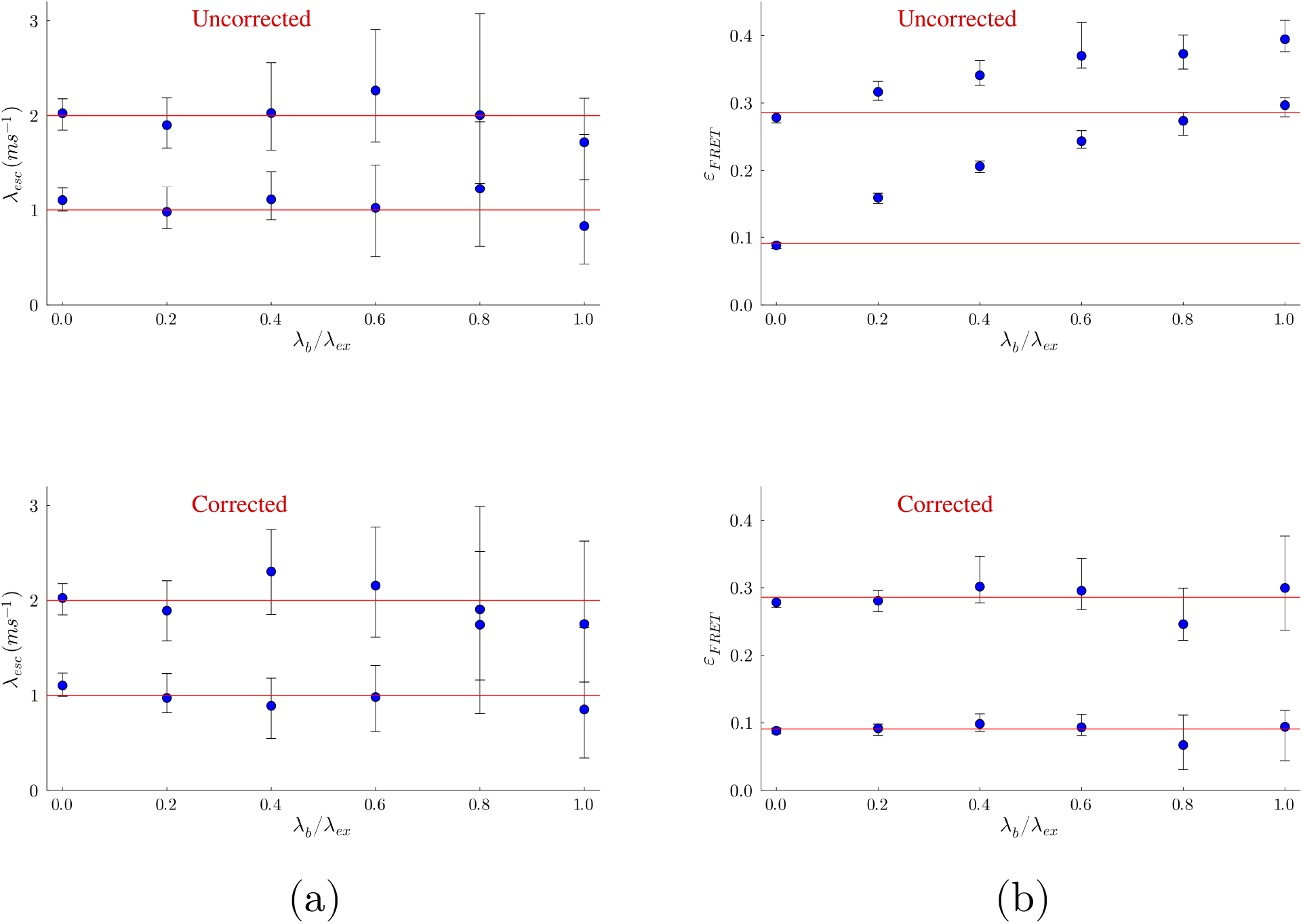
System transition rates λ_*esc*_ and FRET efficiencies *ε_FRET_* as functions of increasing donor and acceptor background fraction λ_*bg*_/λ_*ex*_. To produce these plots, we generated synthetic data with an excitation rate of λ_*ex*_ = 10 ms^-1^, and escape rates λ_*esc*_ = 1 and 2 ms^-1^ for the two system states, respectively, same as Fig. 5, while increasing the fraction of background photons (donor and acceptor) from 0 to 50% (λ_*bg*_/λ_*ex*_ = 1). In each plot, the blue dots represent the median values (50% quantile), and the ends of the attached confidence intervals represent the 5% and 95% quantiles. Furthermore, ground truths are shown with red horizontal lines. In panel (a), we show two plots showing system transition rates estimated by the BNP-FRET sampler when corrected and uncorrected for crosstalk. We show similar plots for FRET efficiencies in panel (b). In all plots, we see that as background is increased, the confidence intervals get bigger (the posterior gets wider) and the estimates become unreliable after λ_*bg*_/λ_*ex*_ > 0.6. Additionally, as expected, if unaccounted for, FRET efficiencies start to merge with increasing background as photons originating from FRET events significantly reduce.

#### 4.1.4 An example with IRF

To demonstrate the effect of the IRF as described in Sec. 2.4.3, we generated new synthetic data for a single fluorophore (with no FRET for simplicity alone) with an escape rate of λ_*d*_ = 2.0 ns^-1^ (similar to that of Cy3 dye [86]) being excited by a continuous-wave laser at a high excitation rate λ_*ex*_ = 0.01 ns^-1^. For simplicity, we approximate the IRF with a truncated Gaussian distribution about 96 ps wide with mean at 48 ps. We again analyze the data with two versions of our method, both incorporating and neglecting the IRF. The results are depicted in Fig. 12, where the posterior is narrower when incorporating the IRF. This is especially helpful when accurate lifetime determination is important in discriminating between different system states. By contrast, accurate determination of lifetimes (which span ns timescales) do not impact the determination of much slower system kinetics from one system state to the next.

**Figure 12:**
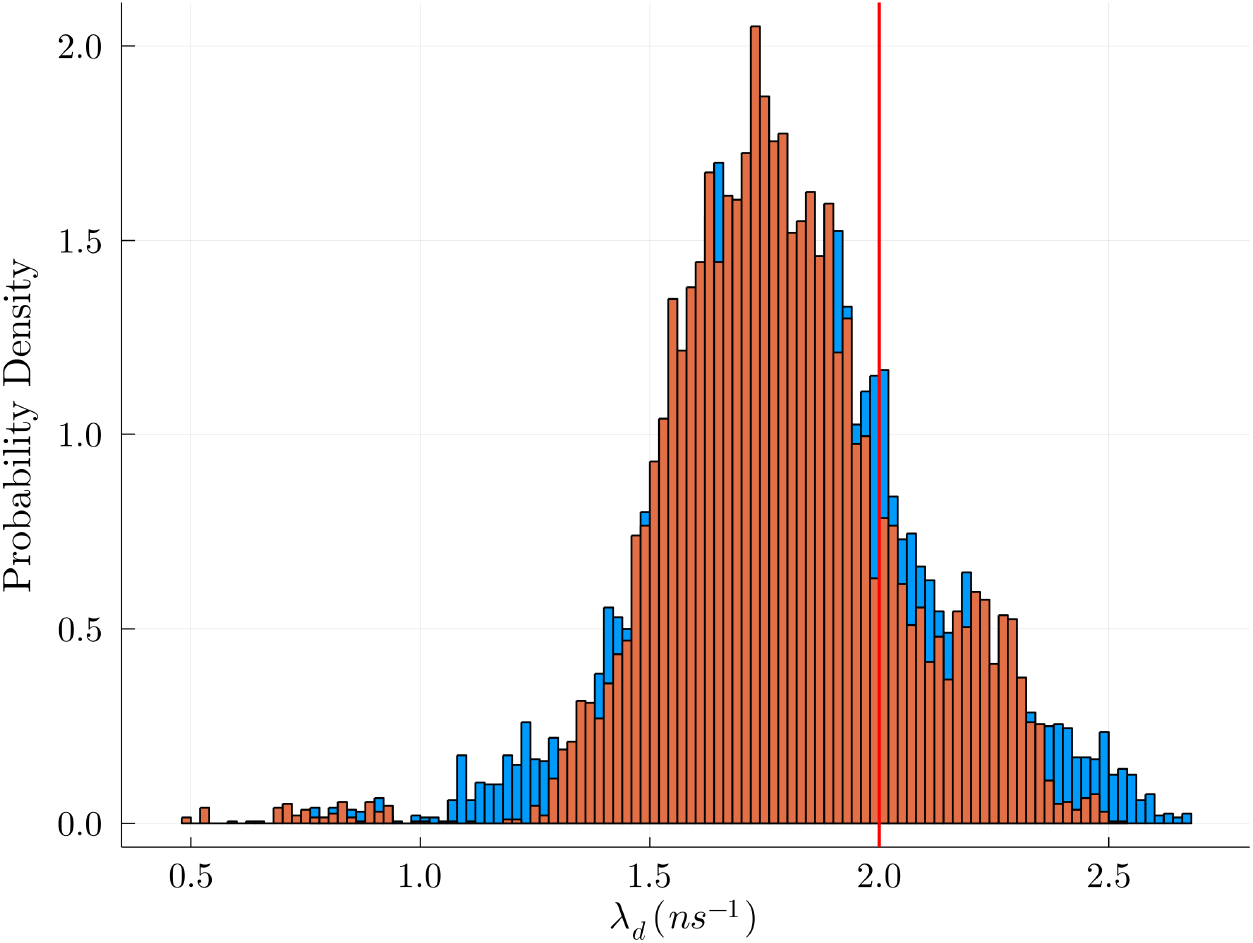
Effects of IRF. Both histograms show the fluorophore’s inverse lifetime with ground truth shown by a red line. The bias in the peaks away from the ground truth arises from the limited amounts of data being used to learn the posterior shown. Corrected model (orange) reduces the histogram’s breadth as compared with the uncorrected model (blue). We conclude from the small effects of correcting for the IRF that, predictably, the IRF may be less important under continuous illumination. By contrast, under pulsed illumination to be explored in the third companion manuscript [64], the IRF will play a more significant role.

### 4.2 A Nonparametric Example

Here, we demonstrate our method in learning the number of system states by analyzing approximately 600 ms (≈120000 photons) of synthetic smFRET time trace data with three system states under pulsed illumination with 25 ns interpulse window; see Fig. 13. This example utilizes the iHMM method described in Sec. 3.2.2 and discussed in greater depth in the third companion manuscript [64]. Using realistic values from the third companion manuscript [64], we set the excitation probability per pulse to be 0.005. Furthermore, kinetics are set at 1.2 ms escape rates for the highest and lowest FRET system states, and an escape rate of 2.4 ms for an intermediate system state. Our BNP method simultaneously recovered the correct system state transition probabilities and thereby the number of system states along with other parameters, including donor and acceptor relaxation rates and the per-pulse excitation probability. By comparison, a parametric version of the same method must assume a fixed number of system states *a priori.* Assuming, say, two system states results in both higher-FRET system states being combined together into one system state with a FRET efficiency of 0.63 and a lifetime of about 0.6 ms; see Fig. 13(c).

**Figure 13:**
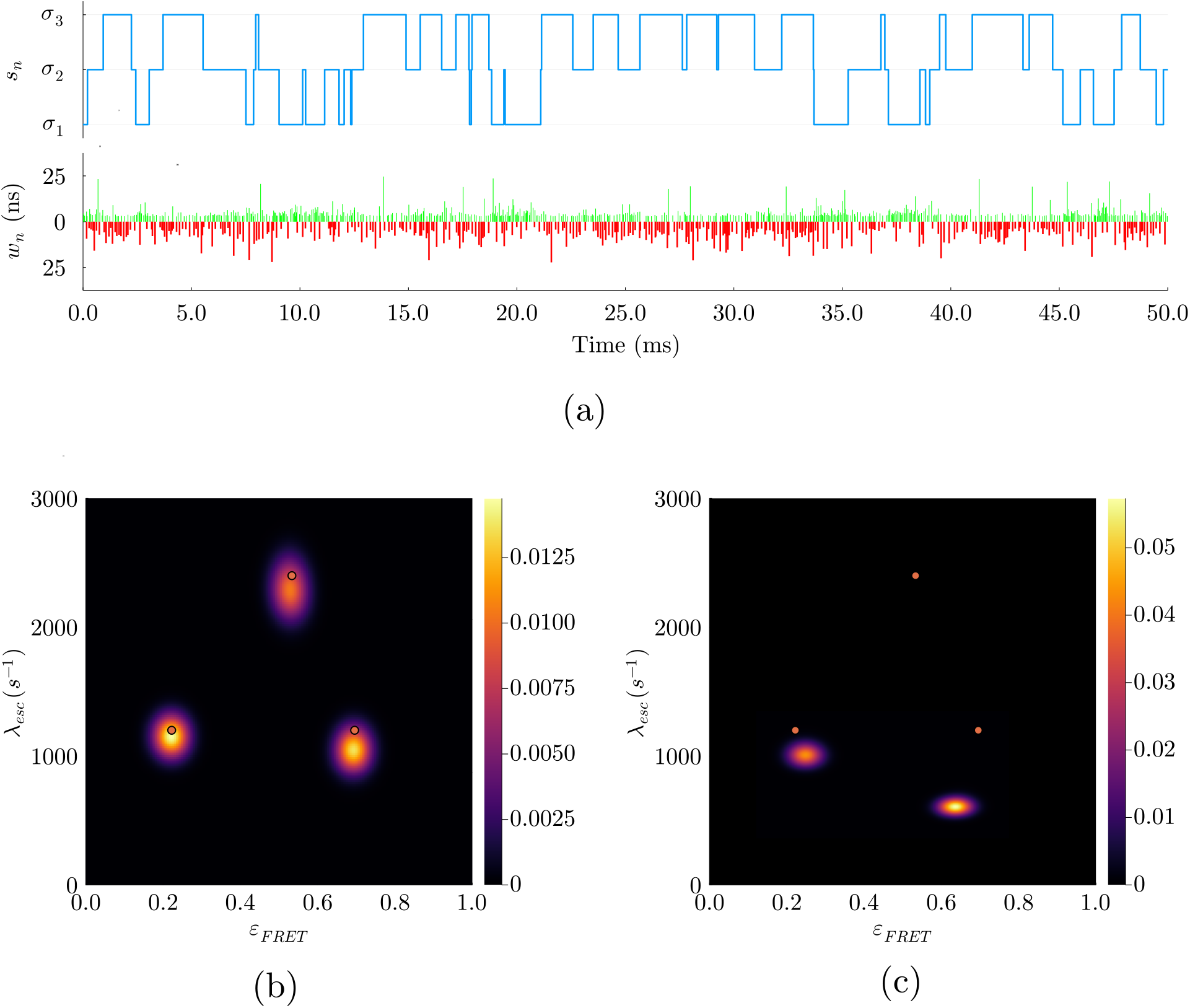
Demonstration of nonparametric analysis on synthetic pulsed data. In (a) we show simulated data for a pulsed illumination experiment. It illustrates a trajectory with three system states labeled *s_n_* in blue and the corresponding photon arrivals with the vertical length of the red or green line denoting the lifetime observed in nanoseconds. Here (b) and (c) show the bivariate posterior for the escape rates λ_*esc*_ and FRET efficiencies *ε_FRET_*. The ground truth is again shown with red dots. The learned posterior using the two system state parametric model (panel (c)) combines high FRET states into one averaged system state with a long lifetime. The nonparametric model correctly infers three system states, as shown in (b).

## 5 Discussion

In this paper, we have presented a complete framework to analyze single photon smFRET data which includes a photon-by-photon likelihood, detector effects, fluorophore characteristics, and different illumination methods. We demonstrated how modern Bayesian methods can be used to obtain full distributions over the parameters, and discussed limitations posed by the photon budget and excitation rate. Additionally, we have shown how to implement a nonparametric inverse strategy to learn an unknown number of system states.

Our method readily accommodates details relevant to specialized smFRET applications. For instance, we can analyze spatial and temporal dependence of excitation by simple modification of generator matrices included in Eq. 21. This is useful in experiments employing pulsed illumination, (Sec. 2.5), as well as alternating-laser excitation (ALEX). In particular, ALEX is used to directly excite the acceptor label, either as a way to gain qualitative information about the sample [63, 87], to reduce photobleaching [63, 87], or to study inter-molecular interactions [16]. Similarly, the generator matrix in Eq. 21 can easily be expanded to include any number of labels, extending our method beyond two colors. Three color smFRET experiments have revealed simultaneous interactions between three proteins [88], monitored conformational subpopulations of molecules [89], and improved our understanding of protein folding and interactions [2, 16, 90–92].

As the likelihood (Eq. 21) involves as many matrix exponentials as detected photons, the computational cost of our method scales approximately linearly with the number of photons and quadratically with the number of system states. For instance, it took 5 hours to analyze the data used to generate Fig. 5 on a regular desktop computer. Additions to our model which increase computational cost include: 1) IRF; 2) pulsed illumination; and 3) BNPs. The computational cost associated to the IRF is attributed to the integral required (see Sec. 2.4.3). The cost of the likelihood computation in the pulsed illumination case depends linearly with pulse number, rather than with photon number (see Eq. 51). This greatly increases the computational cost in cases where photon detections are infrequent. Finally, BNPs necessarily expand the dimensions of the generator matrix whose exponentiation is required (Eq. 21) resulting in longer burn-in time for our MCMC chains.

As a result, we have optimized the computational cost with respect to the physical conditions of the system being studied. First, inclusion of the IRF can be parallelized, potentially reducing the time-cost to a calculation over a single data acquisition period. In our third companion manuscript [64], dealing with pulsed illumination, we improve computational cost by making the assumption that fluorophore relaxation occurs within the window between consecutive pulses, thereby reducing our second order structure herein to a first order HMM, and allowing for faster computation of the likelihood in pulses where no photon is detected. In [64], we mitigate the computational cost of (3) by assuming physically motivated timescale separation.

As it stands, our framework applies to smFRET experiments on immobilized molecules. However, it is often the case that molecules labeled with FRET pairs are allowed to diffuse freely through a confocal volume, such as in the study of binding and unbinding events [17], protein-protein interactions [17, 63], and unhindered conformational dynamics of freely diffusing proteins [63, 93]. Photon-by-photon analysis of such data is often based on correlation methods which suffer from bulk averaging [40, 42, 43]. We believe our framework has the potential to extend Refs. [50, 73, 94] to learn both the kinetics and diffusion coefficients of single molecules.

Additionally, our current framework is restricted to models with discrete system states. However, smFRET can also be used to study systems that are better modeled as continuous, such as intrinsically disordered proteins which include continuous changes not always well approximated by discrete system states [17, 95]. Adopting an adaptation of Ref. [54] should allow us to generalize this framework and instead infer energy landscapes, perhaps relevant to protein folding [96, 97], continuum ratchets as applied to motor protein kinetics [98], and the stress modified potential landscapes of mechano-sensitive molecules [99].

To conclude, we have presented a general framework and demonstrated the importance of incorporating various features into the likelihood while learning full distributions over all unknowns including system states. In the following two companion manuscripts [64, 78], we specialize our method, and computational scheme, to continuous and pulsed illumination. We then apply our method to interactions of the intrinsically disordered proteins NCBD and ACTR [78] under continuous illumination, and the kinetics of the Holliday junction under pulsed illumination [64].

## 6 Code availability

The BNP-FRET software package is available on Github at https://github.com/LabPresse/BNP-FRET

## 8 Acknowledgments

We thank Weiqing Xu and Dr. Zeliha Kilic for regular feedback and help, especially during the development of the nonparametrics samplers, and Dr. Irina Gopich for discussions. S. P. acknowledges support from the NIH NIGMS (R01GM130745) for supporting early efforts in nonparametrics and NIH NIGMS (R01GM134426) for supporting single photon efforts. Bulk of the computations were performed on Agave and Sol supercomputers at ASU.

